# Therapeutic delivery of albumin-binding siRNA targeting IRS2 to diverse cell types reduces mammary tumor growth

**DOI:** 10.1101/2025.07.29.667434

**Authors:** Claire E Tocheny, Julianna E Buchwald, Christopher D Dahlke, Hassan H Fakih, Jennifer S Morgan, Ashley Summers, Christi A Wisniewski, Samuel O Jackson, Ji-Sun Lee, Michael-Anthony Card, Dimas Echeverria, Cornelia Peterson, Arthur M Mercurio, Anastasia Khvorova, Leslie M Shaw

## Abstract

Oligonucleotide therapeutics are a new class of drugs that enable robust and sustained modulation of gene expression. However, achieving efficient delivery of siRNAs to tumors is a challenge for therapy. Here, we demonstrate that fully chemically modified siRNAs conjugated with an albumin-binding dendrimer are efficiently delivered to both neoplastic and stromal/immune cells within primary TNBC mammary tumors. siRNAs were designed to selectively target IRS2, a signaling adaptor of insulin and insulin-like growth factor signaling that has been implicated in aggressive breast cancers. These siRNAs reduced Irs2 expression in tumor and stromal cells without causing hyperglycemia, resulting in reduced tumor growth that was associated with decreased vascularization and alterations in macrophage polarization and the expression of EMT proteins. This work demonstrates that siRNAs can be delivered to neoplastic and specific stromal populations in mammary tumors and that they can effectively and specifically silence a driver of aggressive breast cancer.

## Introduction

Breast cancer remains a major cause of cancer-related morbidity and mortality, particularly due to the limited efficacy of targeted therapies against aggressive subtypes such as triple negative breast cancer (TNBC). One challenge in developing targeted approaches to effectively treat TNBC is the limited ability to inhibit non-enzymatic, intracellular proteins that play a causal role in cancer progression. The insulin receptor substrate (IRS) proteins, which are key mediators of the insulin and insulin-like growth factor (IGF) signaling (IIS) pathway and some integrin and cytokine receptors, exemplify the functional significance of such proteins in cancer (1). IRS proteins are intracellular adaptor proteins that form signaling complexes with downstream effectors following integrin, growth factor, and cytokine receptor activation to facilitate tumor initiation, growth, metastasis, and chemotherapy resistance (2). Among the IRS family, IRS1 and IRS2 are commonly expressed in tumors, although with varying expression patterns and functions, and differing associations with disease outcomes. IRS2 is more highly expressed than IRS1 in invasive breast cancer subtypes, including TNBC, and its expression and activity are associated with poorer clinical outcomes and aggressive tumor behaviors, including invasion, migration, survival and metastasis (3–9). Targeting IRS2 specifically could be a novel intervention to inhibit the IIS pathway in breast tumors, as prior receptor/ligand-targeted approaches have been unsuccessful due to adverse systemic toxicities, such as hyperglycemia, that are caused by disruption of normal metabolic regulation and that lead to positive feedback of insulin signaling in tumors (10–13). Selective inhibition of IRS2, but not IRS1, could maintain metabolic homeostasis while preventing feedback insulin signaling that drives tumor progression. Traditional antibody and small molecule-based therapies, however, are unlikely to selectively inhibit IRS2 function due to its cytoplasmic localization, non-enzymatic activity, and high degree of structural and functional homology with IRS1, and alternative approaches are needed to develop such a disease-modifying therapeutic.

Small interfering RNAs (siRNAs) are a novel class of therapeutics, with seven drugs approved for clinical use (14–20). Once delivered to cells, siRNAs are incorporated into the RNA-induced silencing protein complex (RISC), which then binds to and degrades complementary mRNA transcripts, preventing target protein translation (21). The programmable nature of therapeutic siRNAs enables silencing of virtually any gene target that is important for cancer progression, regardless of protein structure or cellular localization (22). Furthermore, chemical stabilization of siRNAs can extend their duration of action to 6-12 months after a single clinical dose (23), making this drug class especially attractive for antineoplastic therapy. Achieving efficient and sustained delivery of siRNAs to tumors and diverse cell populations of the tumor microenvironment (TME), however, has been a challenge and a limitation for clinical translation (24,25). Studies that explore siRNA delivery and target gene silencing in both neoplastic cells and the TME in immunocompetent breast tumor models are lacking, but necessary, for advancing siRNA therapeutics for breast cancer treatment.

We have previously shown that conjugation of siRNAs to lipophilic moieties or albumin-binding dendrimers facilitates broad biodistribution and delivery (26,27). Emerging evidence suggests that albumin-binding conjugates may outperform lipophilic conjugates in delivering siRNAs to tumors, but this has not yet been evaluated in immunocompetent breast cancer models (28,29). In this study, we evaluate the impact of siRNA conjugate type on delivery efficiency in primary TNBC tumors and show that an albumin-binding dendrimer, but not lipophilic, conjugate achieves significant tumor accumulation. Moreover, albumin-binding conjugates improve siRNA uptake in both neoplastic cells and cells of the TME. We then identify fully chemically modified siRNAs capable of selectively silencing IRS2, but not IRS1, in both human and mouse models. When conjugated to an albumin-binding dendrimer, these siRNAs demonstrated potent IRS2 silencing and significantly suppressed tumor growth in an immunocompetent model of TNBC. Together, our findings not only establish a strategy to co-target diverse cell populations within tumors using siRNA therapeutics, but also validate IRS2 as a clinically relevant, previously untargetable, intracellular adaptor protein in breast cancer.

## Results

### Albumin-binding dendrimer conjugation improves siRNA delivery to mammary tumors

A barrier to the use of siRNA therapeutics in cancer is delivery of oligonucleotides to tumors and their diverse cell types. Previous studies have shown that lipophilic and albumin-binding conjugates improve functional siRNA delivery to extrahepatic tissues upon systemic administration, which may allow for improved uptake in solid malignancies (26). TNBC and other primary breast tumors are localized within the adipocyte-rich environment of the healthy mammary tissue, which may further promote internalization of lipid-conjugated siRNAs. Emerging evidence also suggests that conjugates that bind to albumin, which has been used as a carrier for existing breast cancer therapies (30), increase plasma circulation time of siRNAs (29,31), a property that may enhance tumor distribution. Thus, we compared TNBC mammary tumor delivery and accumulation of siRNAs conjugated to the 22-carbon fatty acid docosanoic acid (DCA) and an amphiphilic, albumin-binding dendrimer structure (Figure 1A) (26,27).

**Figure 1.**
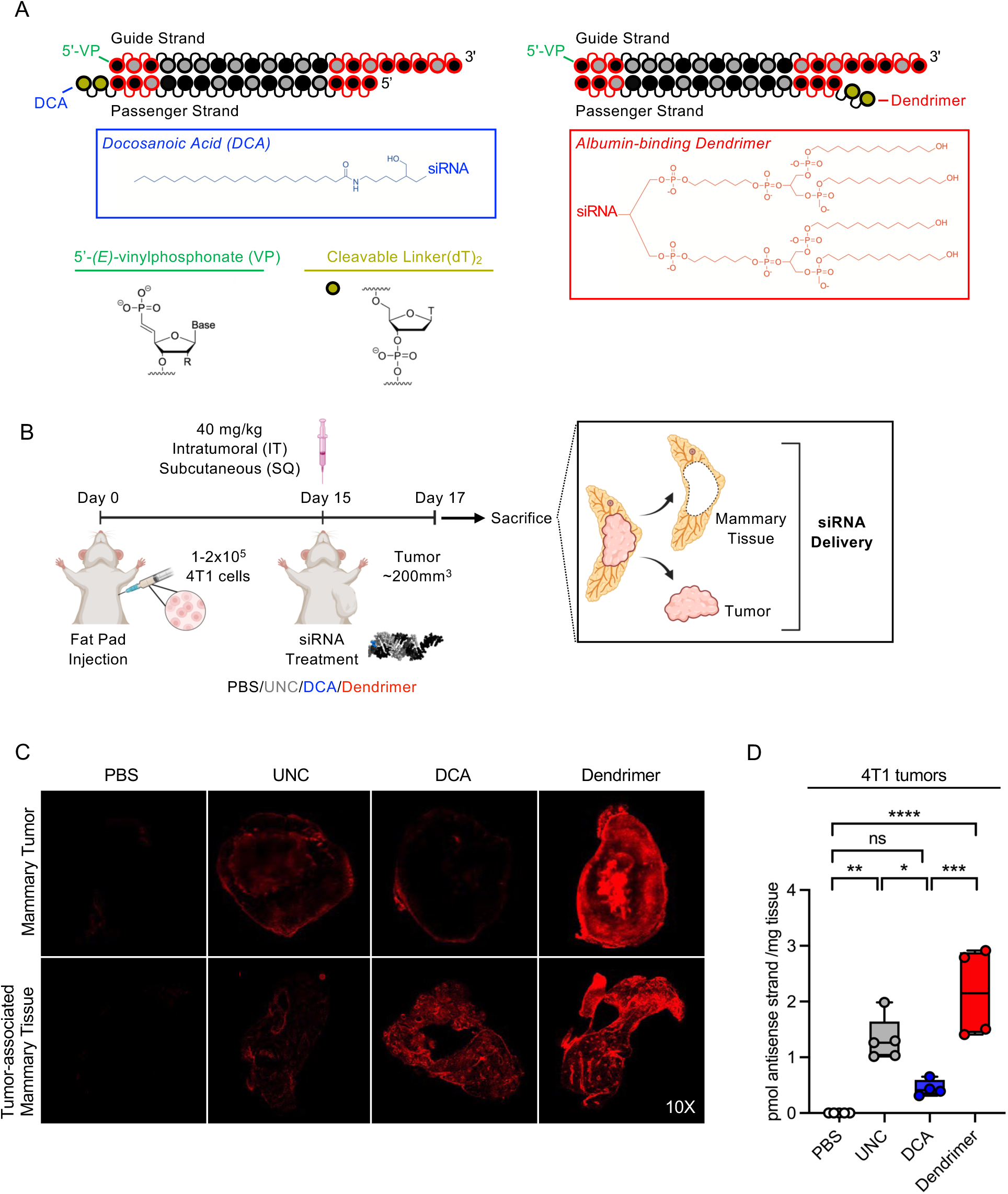
An albumin-binding dendrimer conjugate improves siRNA delivery to mammary tumors. (A) Chemical structures of lipophilic docosanoic acid (DCA), and albumin-binding dendrimer conjugates. (B) 4T1 cells were orthotopically implanted into the fat pads of syngeneic BALB/c mice and mice were treated subcutaneously or intratumorally once with 40 mg/kg conjugated siRNA. (C) Representative whole-tissue fluorescence microscopy images of mammary tumors and tumor-associated mammary fat pad tissue from mice subcutaneously injected with 40mg/kg Cy3-labeled unconjugated, DCA-conjugated, or dendrimer-conjugated siRNAs targeting *Htt* (n=5 mice/group). (D) Tumor accumulation levels of Cy3-labeled siRNA 48 hours after subcutaneous siRNA administration (n=4-5 mice/group). UNC: unconjugated. Line in box-and-whisker plots represents median value, while whiskers represent minimum-to-maximum values. ns = nonsignificant *p<0.05 **p<0.01***p<0.001****p<0.0001 by one-way ANOVA.

Syngeneic mice were injected with 4T1 TNBC tumor cells orthotopically into the mammary fat pad and palpated daily until tumor volumes were approximately 200mm^3^ (Figure 1B). Mice were then injected locally (intratumoral) and systemically (subcutaneous) with Cy3-labeled unconjugated, DCA-conjugated, or dendrimer-conjugated siRNAs targeting the ubiquitously expressed, previously validated housekeeping gene *Htt* (*32*) to assess spatial distribution within tumors. Tumors and adjacent mammary tissue were collected 48 hours post-injection to assess siRNA biodistribution by fluorescence microscopy (Figure 1C). Cy3 fluorescence was largely confined to the site of injection following intratumoral injection, suggesting minimal tumor biodistribution (Supplemental Figure 1). The fluorescence signal was high in tumor-associated mammary tissue, but largely undetectable in the mammary tumors, from mice systemically injected with DCA-siRNAs. In contrast, both tumors and mammary tissue from mice treated with dendrimer-siRNA showed robust, uniform Cy3 fluorescence. Antisense strand concentrations in tumor biopsies demonstrated a ∼4.8-fold increase in dendrimer-siRNA delivery compared to DCA-siRNA (p=0.0004) (Figure 1D), suggesting that conjugation with the albumin-binding dendrimer, but not lipophilic DCA, significantly improves siRNA delivery to mammary tumors.

Tumors are a heterogenous microenvironment of primary neoplastic cells, immune cells, and stromal cells, and target gene silencing in one or more of these cellular compartments may impact tumor biology. To assess the cell-type specificity of dendrimer-siRNA uptake in primary mammary tumors, mice with orthotopic 4T1 tumors were administered PBS or 40mg/kg Cy3-labeled unconjugated or dendrimer-siRNAs when tumors were ∼200mm^3^. A second injection was given 72 hours later, and mice were sacrificed after an additional 72 hours, at which time single-cell suspensions were prepared for multi-color flow cytometry analysis (Figure 2A). We have previously demonstrated that the presence of a Cy3 fluorophore does not significantly alter *in vivo* distribution of lipophilic siRNAs and that quantification of siRNA distribution based on Cy3 fluorescence is consistent with tissue antisense strand accumulation profiles, validating the use of Cy3 fluorescence as a proxy for siRNA delivery (26). Primary tumor cells, defined by high expression of EpCAM, mammary duct epithelium, non-tumor stromal cells, and both adaptive and innate immune (CD45+) cell populations displayed varying degrees of Cy3 fluorescent signal (Figure 2B, gating strategy in Supplemental Figure 2). M1-like macrophage populations, defined by high expression of F4/80, CD11c, and MHC class II, and M2-like macrophages, defined by high expression of F4/80 and CD206, had greater siRNA uptake than T cells, dendritic cells, and other immune populations. Cy3 fluorescence within F4/80 positive macrophages was also confirmed by immunofluorescence staining (Figure 2C). Dendrimer conjugation enhanced siRNA delivery to all identified cell populations, notably a ∼3.8-fold increase in tumor cells (p=0.0006), ∼4-fold increase in non-immune stromal cells (p=0.0002), ∼2.5-5 -fold overall increase in M1-like and M2-like macrophage populations, and ∼2.5-fold increase in dendritic cells (p= 0.0019) and multiple T cell populations compared to unconjugated siRNA (Figure 2D). The percentage of cells in tumors that were Cy3 fluorescent were also significantly increased in most populations after treatment with dendrimer-siRNA compared to unconjugated siRNA, suggesting efficient cellular uptake (Figure 2E). Taken together, our results demonstrate that the albumin-binding dendrimer enhances siRNA delivery to both neoplastic cells and TME populations upon systemic injection, enabling co-targeting to diverse cell types within TNBC tumors.

**Figure 2.**
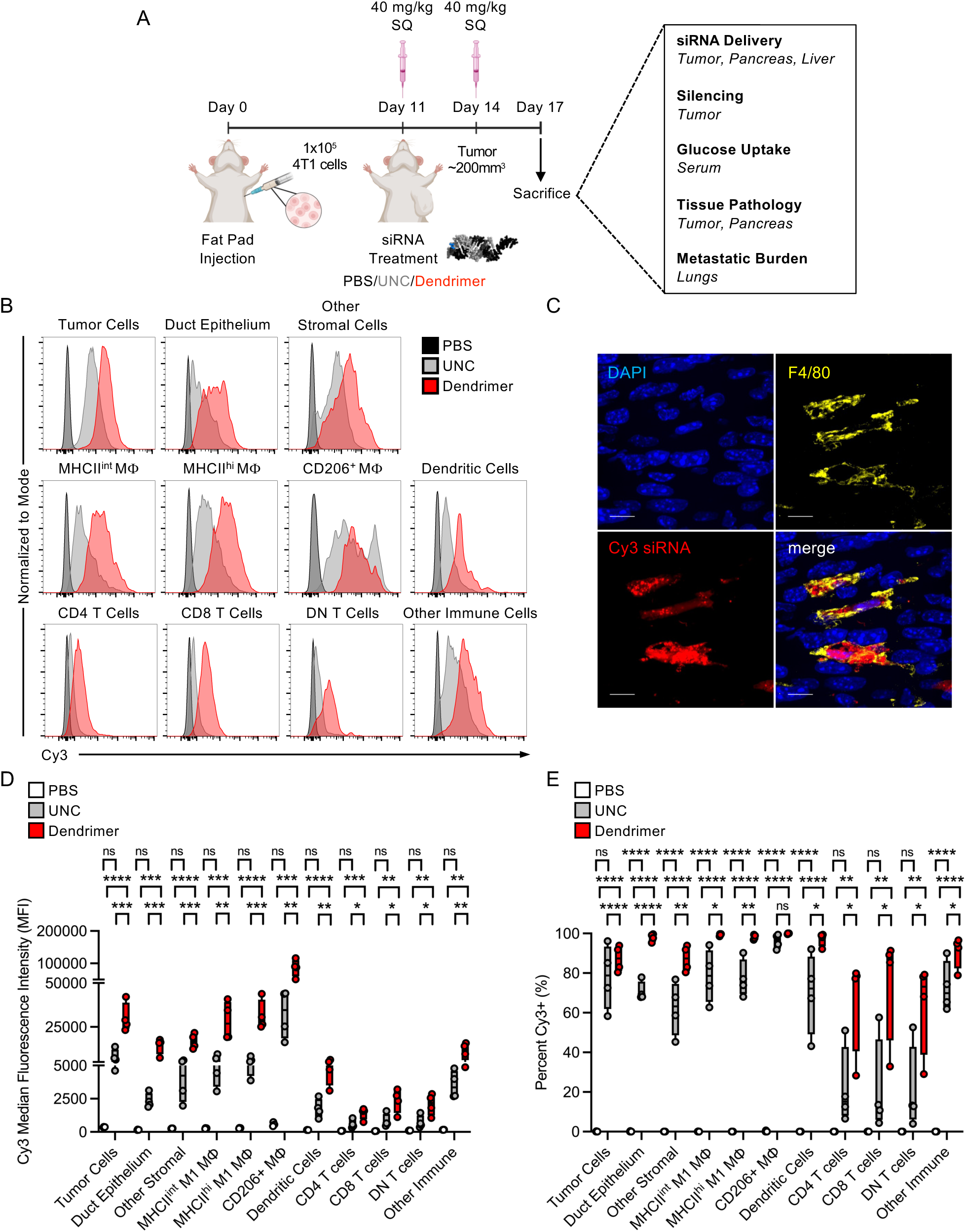
Dendrimer-conjugated siRNA is internalized by tumor and stromal cell types within mammary tumors. (A) 4T1 tumor-bearing BALB/c mice were injected twice with 40 mg/kg Cy3-labeled unconjugated or dendrimer-conjugated siRNA across 6 days to determine cell-specific siRNA delivery in mammary tumors. (B) Representative histograms of Cy3 median fluorescence intensity (MFI) in mammary tumor cell populations. (C) Representative immunofluorescence images of tumor-associated macrophages after siRNA administration. (D-E) Quantification of the Cy3 MFI (D) and the proportion of cell populations with Cy3 fluorescence (E). Analysis of n=4 tumors/group; UNC: unconjugated. MΦ: macrophage, DN: double negative. Line in box-and-whisker plots represents median value, while whiskers represent minimum-to-maximum values. ns = nonsignificant *p<0.05**p<0.01***p<0.001****p<0.0001 by one-way ANOVA (per cell type).

### Identification of human-specific, mouse-specific, and human/mouse cross reactive siRNAs that selectively silence IRS2

Fully chemically modified siRNAs target mRNA transcripts in a highly sensitive and sequence-specific manner, which creates an opportunity to identify therapeutic compounds that selectively silence IRS2. IRS2 is expressed in both primary tumor cells and in cells of the TME and has distinct functions within each compartment that could promote TNBC growth and progression. Assessment of the mechanistic significance of IRS2 within tumor cells or the TME in isolation using immunodeficient mouse tumor models, as well as simultaneously in both compartments using syngeneic mouse models, requires the design of both species-specific and species cross-targeting siRNAs, respectively. We screened a panel of 48 siRNA sequences that were predicted by a proprietary algorithm (33) to selectively silence *IRS2/Irs2* (14 human, 14 mouse, and 20 human/mouse cross-reactive) and were specific to the open reading frame or 3’ untranslated regions (UTR) of *IRS2/Irs2* mRNA transcripts (Figure 3A, Supplemental Table 1). The 3’ end of the siRNA sense strands were conjugated to cholesterol to enable passive uptake for *in vitro* screening (Figure 3B). Screening in two TNBC cell lines, human MDA-MB-231 and mouse 4T1 cells, identified human-specific (si5967), human/mouse cross-targeting (si6530), and mouse-specific (si6548) siRNAs that silenced *IRS2/Irs2* mRNA expression by approximately 50% (Supplemental Figure 3A). The relative median inhibitory concentration (IC50) values were then determined for top siRNA sequences using six-point dose-response studies (Figure 3C-D). Predicted mouse-specific siRNAs did not silence *IRS2* mRNA expression in MDA-MB-231 cells and the predicted human-specific siRNA did not silence *Irs2* mRNA expression in 4T1 cells (Supplemental Figure 3A). We then quantified IRS2 protein in both MDA-MB-231 and 4T1 cells and found that IRS2 protein silencing was consistent with mRNA silencing (approximately 50% reduction) in MDA-MB-231 cells after treatment with si5967 and si6530, but not si6548 (Figure 3E), and that IRS2 protein was reduced approximately 80% in 4T1 cells after treatment with si6530 and si6548, but not si5967 (Figure 3F). Treatment of si6530 and si6548 also significantly reduced IRS2 protein abundance in another cell line, FL83B mouse hepatocytes, further confirming silencing efficacy (Supplemental Figure 3B). Importantly, treatment of all cell lines with si5967, si6530, and si6548 did not silence IRS1 protein abundance (Supplemental Figure 3C). Thus, systematic screening identified highly potent, fully chemically modified siRNAs that efficiently and selectively silence mouse and/or human IRS2, a key advancement towards *in vivo* modulation of IRS2 expression in mammary tumors.

**Figure 3.**
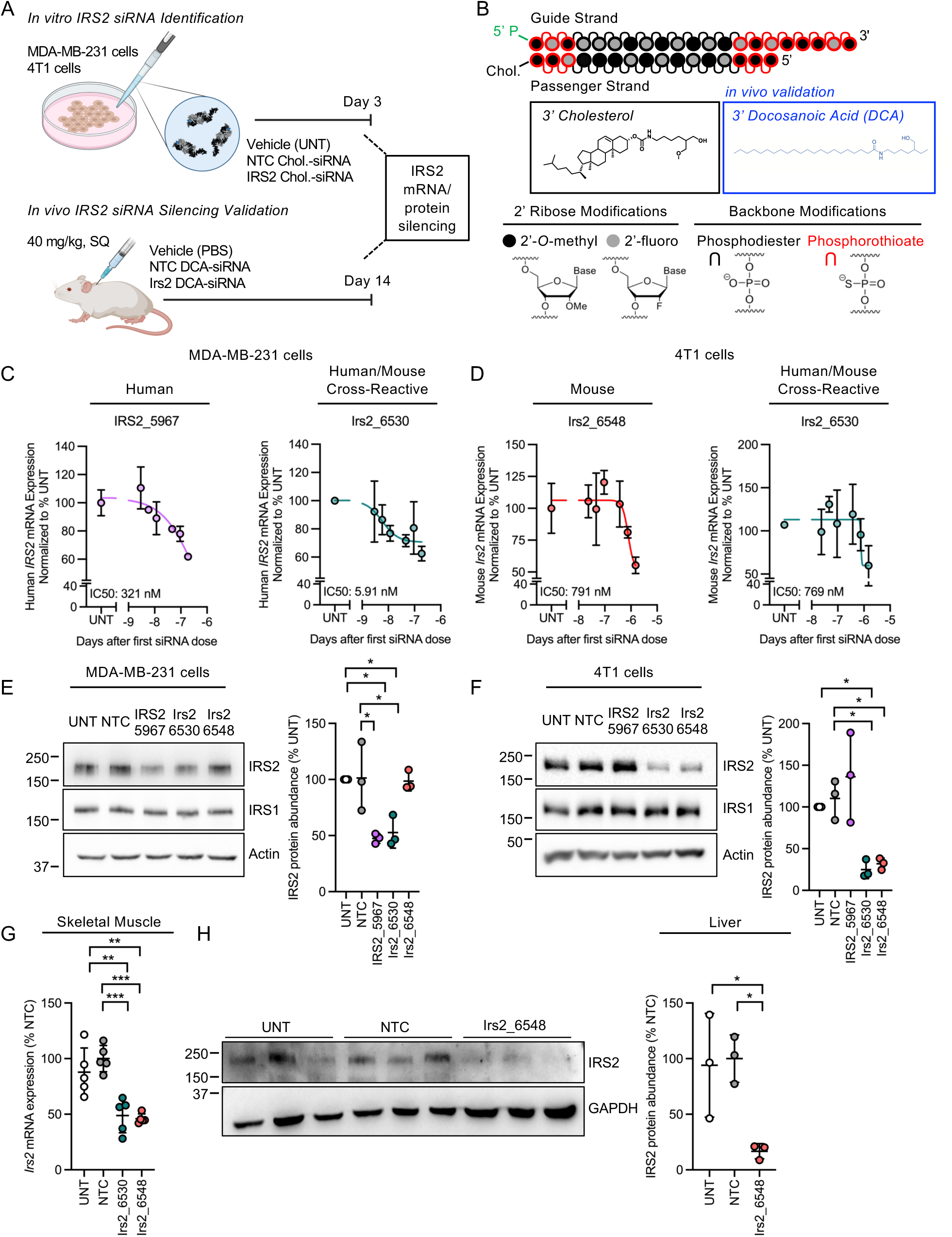
Identification of IRS2-selective siRNAs. (A) *In vitro* and *in vivo* screening of siRNAs in human MDA-MB-231 and mouse 4T1 cells. Cells were treated with fully chemically modified siRNAs conjugated to cholesterol at 0.1875 μM (MDA-MB-231) and 1.5 μM (4T1) for 72 hours prior to IRS2 expression analysis (n=3 independent samples). Mice were subcutaneously injected with 40 mg/kg of the top siRNA candidates and sacrificed 2 weeks post-injection to confirm *in vivo* silencing efficacy . (B) Backbone chemical modifications and conjugate chemical structures of fully chemically modified siRNAs used in *in vitro* and *in vivo* screening. (C-D) Six-point dose-response curves of lead siRNAs 5967 and 6530 in MDA-MB-231 cells (C) and siRNAs 6548 and 6530 in 4T1 cells (D) (n=3 independent samples). (E-F) IRS2 protein abundance in MDA-MB-231 cells (E) and 4T1 cells (F) after 72 hours siRNA treatment (n=3 independent experiments). (G) *Irs2* mRNA expression in skeletal muscle and (H) IRS2 protein abundance in liver 2 weeks after treatment with PBS, NTC DCA-siRNA, or IRS2 DCA-siRNA (n= 3-5 mice/group). UNT: untreated control, NTC: nontargeting control. Data presented as means ±SD. *p<0.05 **p<0.01 ***p<0.001 by one-way ANOVA.

We then assessed *in vivo* silencing efficacy of si6530 and si6548. Naïve mice were subcutaneously injected with si6530 or si6548 (40 mg/kg in 150 μL PBS), and silencing was evaluated after 2 weeks. The 3’ end of the siRNA sense strands were conjugated to DCA to enable broad tissue distribution, while 5’ (*E*)-vinylphosphonate and 3’ extended nucleic acid (ExNA) modifications were added to the siRNA antisense strands to improve 5’ phosphate stability and limit exonuclease-mediated 3’ trimming, respectively (34). DCA is a well-validated conjugate that provides broad distribution to many tissues, including muscle and liver (26,35), where IRS2 is expressed. Use of DCA-conjugated siRNAs, therefore, enabled validation of *in vivo* efficacy before advancing our identified compounds towards more complex tumor models. Treatment with si6530 and si6548 led to 50% reduction in *Irs2* mRNA expression in skeletal muscle (Figure 3G) (26,35), but protein expression was low and difficult to detect by immunoblot. Therefore, IRS2 protein silencing was confirmed in the liver that expresses high levels of IRS2. IRS2 expression was suppressed by approximately 75% by the mouse-specific si6548 (Figure 3H). We next confirmed robust, dose-dependent IRS2 protein silencing after treatment with si6548 in 4T1 cells, the intended cell line for our *in vivo* tumor studies, by performing four-point dose response studies (Supplemental Figure 3D). Given these results, we selected si6548 (hereafter referred to as *siIrs2*) as our top candidate for subsequent *in vivo* syngeneic tumor studies.

### siRNA-mediated silencing of *Irs2* reduces mammary tumor growth

Previous studies have implicated IRS2 in multiple aspects of breast cancer progression, including tumor initiation and stemness, tumor cell migration and invasion, and metastasis using *Irs2* knockout transgenic and orthotopic mouse tumor models (6,8,9,36). We determined that the albumin-binding dendrimer conjugate enables superior siRNA delivery to mammary tumors compared to lipophilic DCA (Figure 1C-D). Thus, to evaluate the impact of acute siRNA-mediated *Irs2* silencing on mammary tumor progression, we injected Cy3-labeled, dendrimer-conjugated *siIrs2* and a nontargeting control siRNA (*siNTC*) into mice bearing 4T1 tumors following the same siRNA injection protocol as was performed for flow cytometry studies (Figure 2A). A second cohort of tumor-bearing mice was also injected with a dendrimer-siRNA previously validated to potently silence the inflammatory mediator JAK1 (*siJak1*) as an additional control for non-specific effects of siRNA sequence on tumor silencing (*37*). Tumors exhibited a slower growth rate in mice treated with *siIrs2* with significant reductions (p=0.0008) demonstrated following administration of the second siRNA dose (Figure 4A, Supplemental Figure 4A). As a result, terminal tumor volumes at sacrifice were significantly smaller (22.9% reduction, p=0.0066) in mice treated with *siIrs2* than those from mice treated with *siNTC* (Figure 4B). While significant differences in tumor growth rate were observed in mice after administration with the second siRNA dose of *siJak1*, no differences in terminal tumor volumes were observed (Supplemental Figure 4B-D). Robust siRNA delivery to tumors was first established by fluorescence microscopy (Figure 4C, Supplemental Figure 4E). We also confirmed antisense strand accumulation in tumor biopsies in a parallel assay that is independent of Cy3 fluorescence (Figure 4D, Supplemental Figure 4F). Both the center and periphery of tumors contained substantial concentrations of *siIrs2* and *siJak1* antisense strand, suggesting penetration of siRNA throughout tumor tissue. As expected, siRNAs were also robustly delivered to the liver. *Irs2* and *Jak1* mRNA silencing was then assessed in biopsies sampled from non-necrotic areas of mammary tumors. Administration of *siIrs2* and *siJak1* led to significantly reduced *Irs2* (26.7%, p=0.0042) and *Jak1* (20.9%, p=0.0025) mRNA expression, respectively (Figure 4E, Supplemental Figure 4G), indicating successful silencing of two different target genes in a sequence-specific manner. Histological analysis of tumors from mice treated with either *siNTC* or *siIrs2* revealed a trend towards decreased mitotic counts in tumors from mice treated with *siIrs2* compared to *siNTC* (p=0.0534), and no difference in the percentage of necrosis between groups (Supplemental Figure 5A-B). Together, these findings confirm that albumin-binding siRNA is robustly delivered to mammary tumors to enable sequence-specific gene silencing and show, importantly, that modulation of IRS2 expression inhibits TNBC tumor growth.

**Figure 4.**
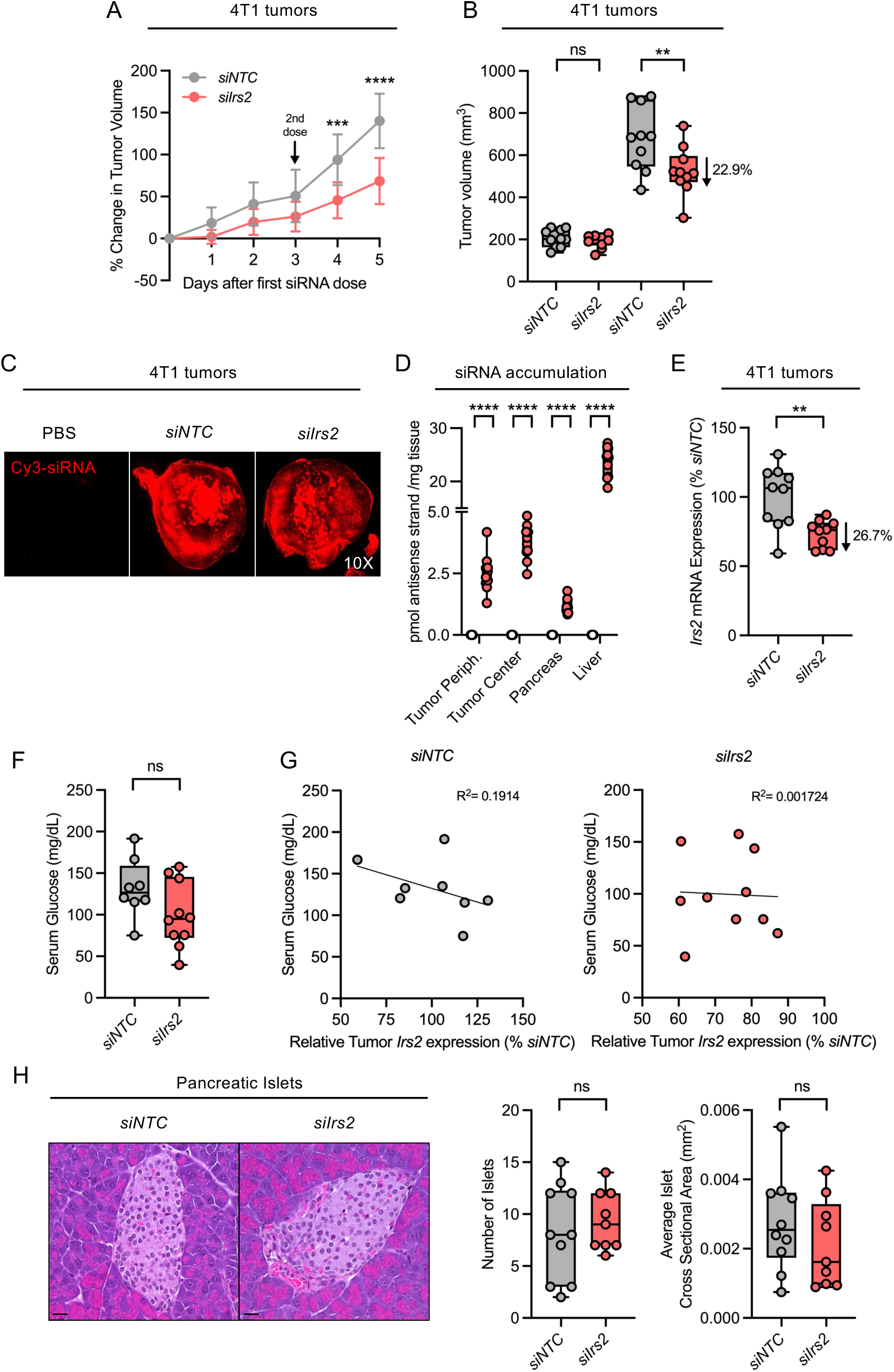
Administration of siRNAs targeting *Irs2* enables tumor gene silencing and inhibits tumor growth with causing hyperglycemia. (A) 4T1 mammary tumor growth rates after *siNTC* or *siIrs2* administration. (B) Tumor volumes before and after *siNTC* or *siIrs2* administration. (C) Representative fluorescence microscopy of 4T1 mammary tumors from BALB/c mice subcutaneously injected with Cy3-labeled dendrimer-conjugated *siNTC* or *siIrs2* or non-targeting control (NTC). (D) Accumulation of Cy3-labeled siRNA targeting *Irs2* in 4T1 mammary tumor, pancreas, and liver tissue from BALB/c after subcutaneous administration of two 40 mg/kg doses across 6 days. (E) *Irs2* mRNA expression in 4T1 mammary tumor biopsies from mice treated with *siNTC* or *siIrs2*. (F) Terminal serum glucose concentrations in 4T1 tumor-bearing mice after treatment with *siNTC* or *siIrs2*. (G) Comparison of serum glucose levels and relative tumor *Irs2* mRNA expression in *siNTC* or *siIrs2*-treated tumor-bearing BALB/c mice. Relative *Irs2* mRNA expression is presented as a percentage of average *Irs2* mRNA expression in NTC siRNA-treated mice. (H) Representative H&E images and quantification of pancreatic islet number and size in *siNTC* or *siIrs2*-treated 4T1 tumor-bearing BALB/c mice. Analysis of n=8-10 mice/group. NTC: nontargeting control. scale bar = 20 μm. Line in box-and-whisker plots represents median value, while whiskers represent minimum-to-maximum values. ns=nonsignificant *p<0.05**p<0.01***p<0.001****p<0.0001 by Student’s t-test (Figures 4B (per tissue type), C, D (per time point), F, H); ns=nonsignificant **p<0.01 by one-way ANOVA (Figure 4E). Analysis of n=8-10 mice/group. R^2^ determined by simple linear regression (Figure 4G).

### Acute treatment with *siIrs2* does not cause hyperglycemia

A major concern regarding therapeutic inhibition of components of the IIS pathway is disruption of insulin-dependent regulation of glucose uptake in metabolic tissues, such as the liver or muscle, resulting in hyperglycemia. Mice lacking *Irs2* expression develop insulin resistance, which leads to hyperglycemia, and exhibit progressive loss of pancreatic islet mass (38). Analysis of terminal serum glucose concentrations revealed that glucose was not elevated in mice treated with *siIrs2* and there were no significant differences in serum glucose concentrations between mice treated with *siIrs2* and *siNTC* (Figure 4F). Importantly, no correlation was observed between *Irs2* tumor mRNA expression and serum glucose levels in siRNA-treated mice (Figure 4G). Moreover, neither the number nor size of pancreatic islets were decreased in mice treated with *siIrs2* compared to *siNTC* (Figure 4H), further supporting the conclusion that acute treatment with *siIrs2* does not significantly perturb systemic glucose homeostasis.

### Suppression of *Irs2* modulates the mammary tumor microenvironment

Suppression of tumor growth by *siIrs2* may be the result of both tumor cell-intrinsic and - extrinsic functions of IRS2, as this adaptor is expressed in primary tumor cells and in some cell populations of the TME. Stromal/immune cells that express IRS2 include macrophages, where it acts as a signaling adapter for the IL-4 receptor to drive polarization to the M2 phenotype, endothelial cells, and fibroblasts (39–41). To examine the mechanism by which *siIrs2* inhibits tumor growth, we first compared tumor immune populations from mice treated with *siIrs2* and *siNTC*, as *Irs2* silencing could alter the composition of the TME. Flow cytometric analysis revealed that the proportion of tumor dendritic cells, total macrophages, T cells, and other, undefined immune populations did not significantly differ between treatment groups (Figure 5A). Although the frequency of total tumor-associated macrophages (TAMs) was similar in *siIrs2*- and *siNTC*-treated mice, the frequency of macrophages that highly expressed CD206 (M2-like) was significantly decreased (p=0.0099) in tumors from mice treated with *siIrs2* compared to *siNTC* (Figure 5B). As a result, the ratio of macrophages that highly expressed CD11c (M1-like) to those that highly expressed CD206 (M2-like) was significantly increased in *siIrs2*-treated mice (p=0.0286) (Figure 5C).

**Figure 5.**
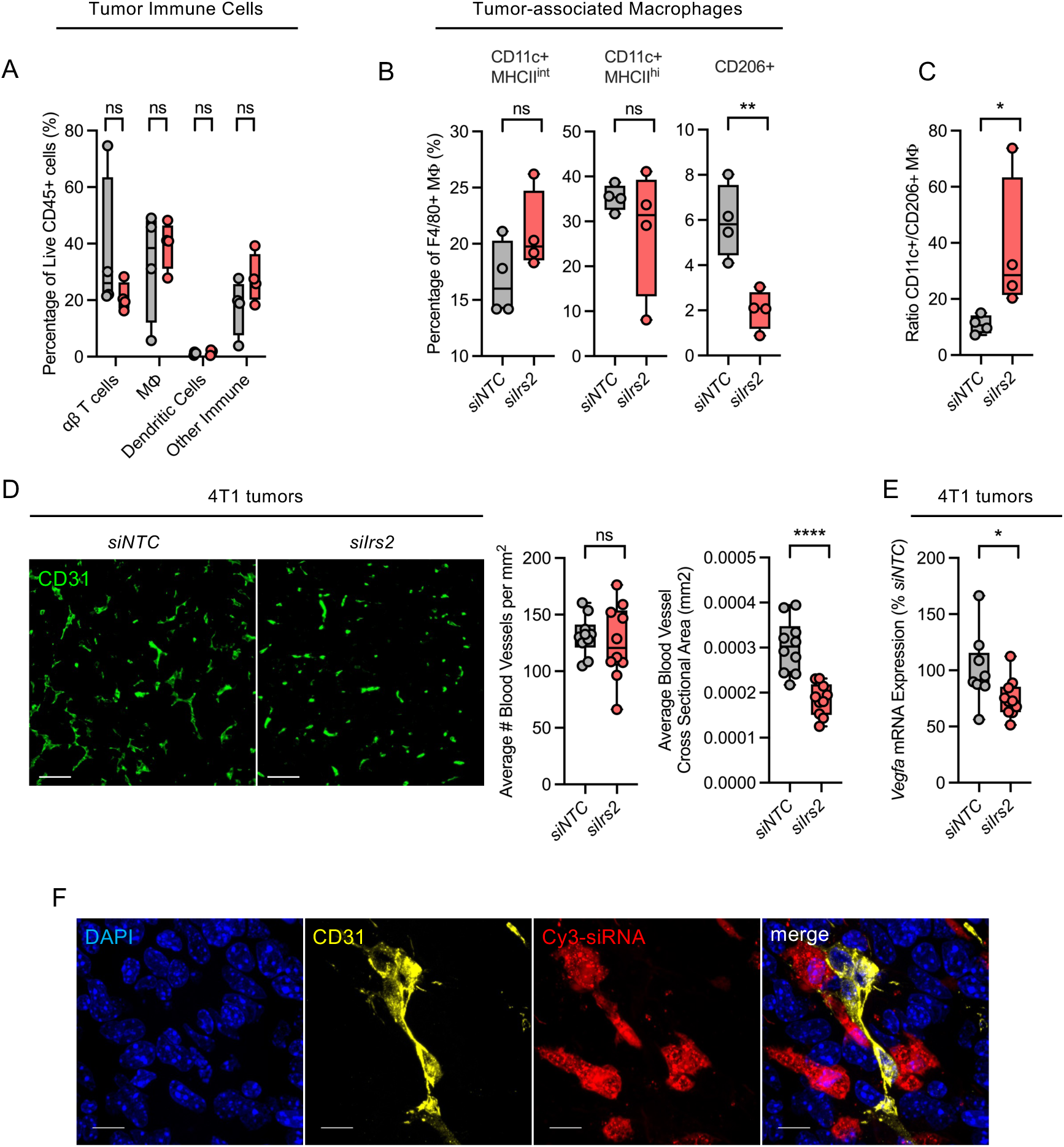
*Irs2* silencing modulates the mammary tumor microenvironment. (A-B) Flow cytometric analysis of the proportion of immune cell subsets (A) and macrophage subsets (B) within 4T1 mammary tumors from BALB/c mice treated with *siNTC* or *siIrs2*. (C) Quantification of the ratio of M1-like to M2-like macrophages in siRNA-treated tumors. (D) Representative immunofluorescence images and quantification of CD31-positive blood vessels in mammary tumors from *siNTC* or *siIrs2*-treated mice. (E) Relative *Vegfa* mRNA expression in tumor biopsies from mice treated with *siNTC* or *siIrs2*. (F) Representative immunofluorescence images of mammary tumor blood vessels stained for CD31 after siRNA administration (Cy3). Analysis of n=10 mice/group (Figure 5D,E) and of n=4 tumors/group (Figure 5A-C, F). NTC: nontargeting control, MΦ: macrophage. scale bars 100 μm (Figure 5A), or 10 μm (Figure 5F). Line in box-and-whisker plots represents median value, while whiskers represent minimum-to-maximum values. *p<0.05 by Mann-Whitney test (Figure 5C) ns nonsignificant *p<0.05**p<0.01****p<0.0001 by Student’s t-test (Figure 5A (per cell type), B, 5D-E).

M2-like TAMs are a key driver of angiogenesis, a process that provides nutrients and oxygen necessary to sustain tumor expansion and spread (42–44). To assess the impact of *Irs2* silencing on tumor vascularization, we measured the density and average area of blood vessels in the mammary tumors using the expression of CD31 as a marker of endothelial cells (Figure 5D). While blood vessel density was similar in tumors across groups, average vessel cross sectional area was significantly decreased (p<0.0001) in tumors from mice treated with *siIrs2* compared to *siNTC* (Figure 5D). Expression of the growth factor *Vegfa* was also significantly decreased in tumor biopsies from *siIrs2-*treated mice (p=0.0397), suggesting a reduction in pro-angiogenic factors within the TME (Figure 5E). To evaluate if the reduction in tumor vascularization could be to the result of *Irs2* silencing directly in endothelial cells, we evaluated Cy3 fluorescence in CD31 positive cells by fluorescence microscopy (Figure 5F). Co-localization of Cy3 and CD31 staining was not observed, revealing that *siIrs2* was not internalized efficiently by endothelial cells.

### Reduction of *Irs2* expression in mammary tumors promotes tumor cell epithelial differentiation

Studies have reported a tumor cell-intrinsic role for IRS2 in tumor cell survival, epithelial-to-mesenchymal transition (EMT), migration, and invasion (4,7–9,45). We observed that neoplastic cells near the peripheral margins of tumors assumed a more fusiform morphology, indicative of dedifferentiation and EMT. To assess the impact of *Irs2* silencing on this phenotype, we evaluated the expression of the cytoskeletal protein vimentin, which is upregulated in cells that have undergone EMT, in tumors from *siIrs2*- and *siNTC*-treated mice. We found that the number of neoplastic cells that expressed vimentin in tumors from mice treated with *siIrs2* was reduced by ∼75% compared to *siNTC* (p<0.0001), and that the intensity of vimentin expression was decreased by approximately 2-fold in these cells (p=0.001) (Figure 6A). We also evaluated the expression of the epithelial marker E-cadherin in a subset of tumors and observed an ∼8-fold increase in the number of tumor cells that expressed E-cadherin (p=0.0017) and a ∼2-fold increase in the intensity of E-cadherin expression in tumors from mice treated with *siIrs2* (p=0.0257) (Supplemental Figure 5C).

**Figure 6.**
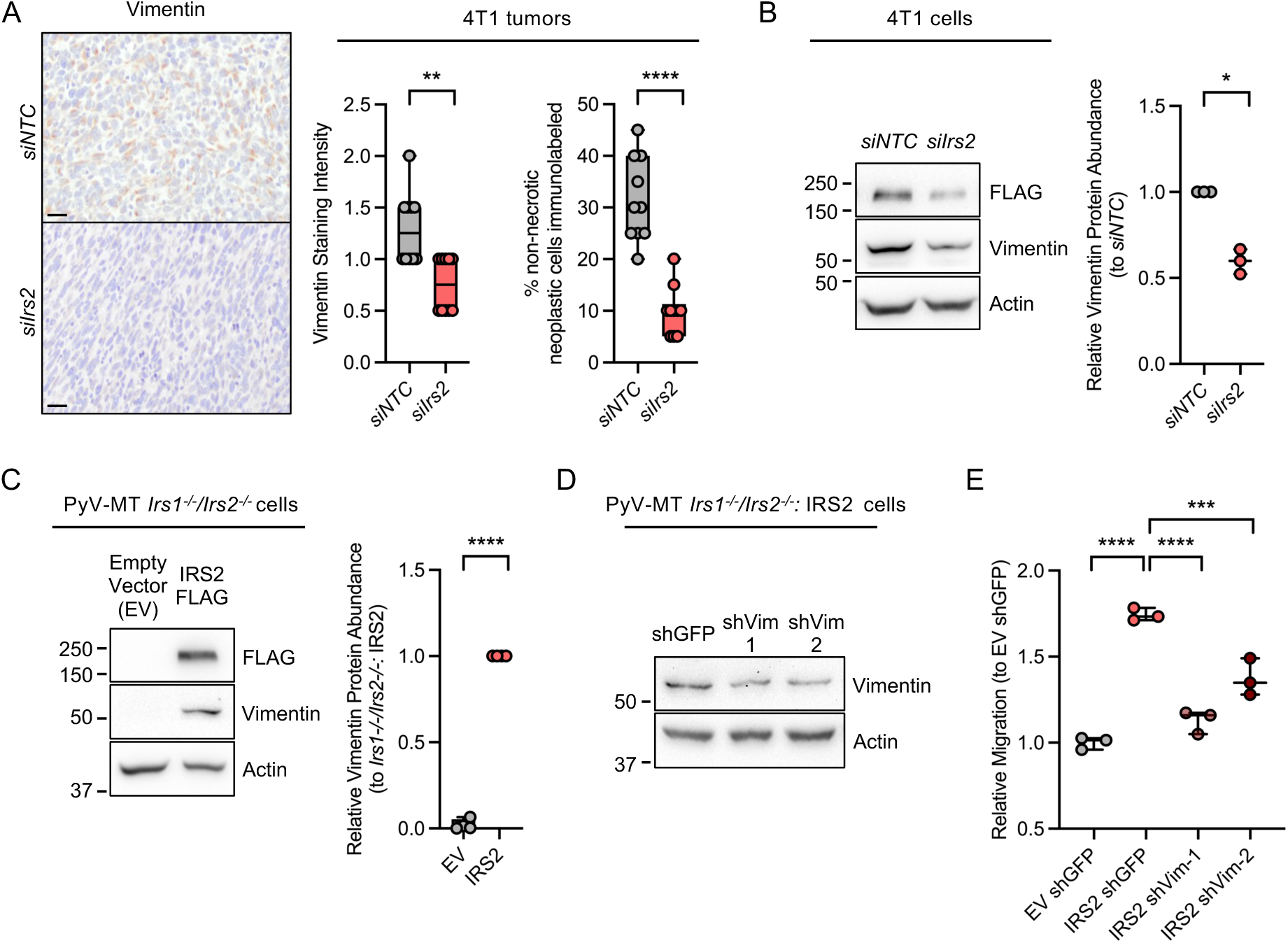
Primary mammary tumors exhibit less mesenchymal features after treatment with *siIrs2*. (A) Representative immunohistochemistry images and quantification of vimentin expression in 4T1 mammary tumors from siRNA-treated mice. (B) Vimentin protein abundance in 4T1 cells 72 hours after *in vitro* treatment with cholesterol-conjugated *siNTC* or *siIrs2.* (C) Vimentin protein abundance in PyV-MT *Irs1^-/-^/Irs2*^-/-^ cells expressing either empty vector (EV) control or IRS2. (D) Representative vimentin protein abundance in PyV-MT *Irs1^-/-^/Irs2*^-/-^: IRS2 cells after treatment with either shRNAs targeting GFP (control) or vimentin. (E) Transwell migration assay of PyV-MT *Irs1^-/-^/Irs2*^-/-^ cells expressing either empty vector (EV) or IRS2 after treatment with shRNAs targeting GFP or vimentin. Analysis of n=10 mice/group (Figure 6A). n=3 (Figure 6B,C,E) independent experiments. Figure 6D is representative of 3 independent experiments. NTC: nontargeting control, EV: empty vector. Scale bars 20μm. Line in box-and-whisker plots (Figure 6A) represents median value, while whiskers represent minimum-to-maximum values. Data presented as means ±SD (Figures 6B,C,E). *p<0.05**p<0.01****p<0.0001 by Student’s t-test (Figures 6A-C) ***p<0.001****p<0.0001 by one-way ANOVA (Figure 6E).

We further dissected the role of IRS2 in regulating vimentin and E-cadherin *in vitro* using 4T1 cells and *Irs1^-/-^/Irs2^-/-^* cells isolated from PyV-MT murine tumors. Treatment of 4T1 cells *in vitro* with cholesterol-conjugated *siIrs2* led to a significant decrease in vimentin protein abundance compared to *siNTC* (p=0.0105), supporting that *in vivo* reduction of vimentin expression was due to tumor cell-intrinsic *Irs2* silencing (Figure 6B). However, E-cadherin protein abundance did not increase in *siIrs2-*treated 4T1 cells *in vitro,* indicating that IRS2 may regulate E-cadherin expression *in vivo* through tumor cell-extrinsic mechanisms (Supplemental Figure 5D). Restoration of IRS2 expression in PyV-MT:*Irs1^-/-^/Irs2^-/-^* cells increased vimentin protein abundance (Figure 6C), suggesting that IRS2 is also sufficient to upregulate vimentin in a tumor cell-intrinsic manner. PyV-MT: *Irs1^-/-^/Irs2^-/-^*cells with restored expression of IRS2 exhibited enhanced migratory potential and knockdown of vimentin expression using 2 different shRNAs in these cells reduced this cell migration (Figure 6D-E). Together our results demonstrate a robust impact of *Irs2* silencing on EMT markers and suggest a shift toward a more differentiated, epithelial phenotype upon suppression of IRS2 expression in mammary tumors.

Tumor cell de-differentiation is connected to breast cancer metastasis(46). Previous studies using knockout transgenic mouse tumor models have found that IRS2 supports breast cancer metastasis (6). We evaluated lung metastatic burden in *siIrs2*- and *siNTC*-treated mice to investigate if acute *Irs2* silencing in mammary tumors impacts dissemination. A similar percentage of mice developed metastases in both groups (8 out of 10 mice), and there was no significant difference in the number of metastatic lesions within the lungs (Supplemental Figure 6A). Although we observed a similar number of micrometastases, defined as lesions with a cross-sectional area less than 0.01mm^2^, in the lungs of mice treated with *siIrs2* and *siNTC*, there were 2-fold fewer metastatic lesions with a cross-sectional area greater than 0.01mm^2^ in mice treated with *siIrs2* compared to *siNTC* (15 vs 30), indicating a downward trend in the size of lung metastases after treatment with *siIrs2* (Supplemental Figure 6B). In summary, our data support that partial modulation of IRS2 expression (25% *Irs2* silencing) has a profound biological impact on both mammary tumor cells and the TME, culminating in a significant reduction in primary tumor growth.

## Discussion

The potential of siRNA therapeutics to offer unprecedented, selective silencing of genes whose products are unable to be targeted by other treatment modalities has led to significant interest in the use of siRNAs as targeted agents in cancer. Achieving efficient delivery to tumors, however, has been a limitation for the clinical translation of this technology in oncology. In this study, we demonstrate that an albumin-binding dendrimer conjugate significantly improves siRNA delivery to TNBC tumors compared to DCA and promotes cellular uptake in tumor cells and heterogenous cell types of the TME. Treatment with dendrimer-siRNAs that selectively silence IRS2, an enzymatically inactive adaptor protein that contributes to TNBC progression, achieves target gene silencing and slows tumor growth without perturbing glucose homeostasis in an immunocompetent model of TNBC. Suppression of *Irs2* led to a decrease in M2-like TAMs and a higher ratio of M1-like to M2-like TAMs, reflecting a shift in macrophage polarization toward a more anti-tumor population. Primary tumors also exhibited diminished and increased expression of the EMT markers vimentin and E-Cadherin, respectfully, and reduced vascularization, reflecting effects on both tumor cells and the TME in response to *Irs2* tumor silencing. Together, our results establish that albumin-binding conjugates enhance functional and cell-specific siRNA delivery in TNBC tumors and provide mechanistic insight into the role of IRS2 in TNBC progression. These findings highlight the feasibility and benefits of siRNA-based inhibition of nonenzymatic proteins in cancer.

Albumin is an attractive carrier to improve the pharmacokinetic properties of siRNA therapeutics in cancer, as it extends the circulation of rapidly cleared drugs and improves drug accumulation at sites of tumor proliferation (47,48). Moreover, albumin is internalized by tumor cells through macropinocytosis and receptor-mediated pathways that promote the intracellular delivery of siRNA (47,49–51). Previous studies have reported that siRNA conjugated to albumin-binding diacyl lipids or nanocomplexes of siRNA and albumin enable enhanced and productive tumor delivery, providing evidence that albumin-binding conjugates may overcome a key barrier in the use of siRNAs as chemotherapeutic agents (25,28,29). These studies, however, did not address the cell-type specificity of siRNA uptake in the heterogeneous TME. Our study reveals additional benefits of albumin-binding siRNA conjugates by demonstrating that dendrimer-siRNA improves cellular uptake in neoplastic cells and diverse stromal and immune cell populations within syngeneic TNBC mammary tumors. These findings demonstrate the ability to co-target multiple cell populations using a single siRNA chemical scaffold with albumin-binding properties. Moreover, our data support that cell-type specific silencing in tumors may also be achieved using albumin-binding siRNA by altering the siRNA antisense strand sequence to target genes that are unique to one cell population.

Our study is the first to demonstrate that acute silencing of IRS2 can impede mammary tumor growth without causing hyperglycemia or reducing pancreatic islet density or size, which obviates a major concern with currently available therapeutics that target the IIS pathway in cancer. Cell-intrinsic roles for IRS2 in regulating tumor cell functions that contribute to disease progression have been previously established using IRS2 knockout tumor models *in vitro* and *in vivo* (6–8,36,45). We observed that acute *in vitro* and *in vivo* siRNA-mediated suppression of *Irs2* reduces expression of vimentin in neoplastic cells, and that tumor cell-intrinsic regulation of this EMT factor by IRS2 impacts migration, a process that underlies invasive tumor growth (52). Treatment with *siIrs2* also alters the TAM compartment and decreases tumor vascularization. Our data suggest that *Irs2* silencing reduces tumor angiogenesis through mechanisms independent of endothelial cell-intrinsic knockdown. IRS2 acts as a signaling adaptor for the IL-4 cytokine receptor to promote M2-like macrophage polarization (41), which generates a pro-tumor, anti-inflammatory macrophage population that promotes tumor growth and progression through the secretion of factors, such as VEGF, that support angiogenesis and metastasis (42). Promotion of tumor vascularization would facilitate tumor cell access to nutrients and oxygen to promote tumor growth. We posit, therefore, that *Irs2* silencing within both tumor cells and macrophages inhibits processes essential for tumor growth, which emphasizes the value of an siRNA approach that co-targets both tumor cells and the TME and the importance of using immunocompetent mouse models to fully evaluate the functional consequences of siRNA targeting. Future studies that provide a more detailed analysis of *Irs2* silencing within tumor cells and specific macrophage subsets will provide a greater understanding of the cell-specific mechanism(s) by which *Irs2* silencing suppresses mammary tumor growth.

A technical challenge in this study is to accurately assess the impact of *Irs2* silencing on tumor metastasis. It is important to note that the 4T1 syngeneic tumor model is particularly aggressive and metastasis occurs very early (53). While our dosing strategy delivered an excess of oligonucleotides to tumor tissue at the time of injection and resulted in a significant reduction in tumor growth, it is likely that metastasis had initiated prior to the first injection of siRNA. Initiation of treatment at earlier stages of TNBC tumor development would address the overall efficacy of IRS2-selective siRNA as a treatment modality for preventing metastatic disease. Moreover, assays that initiate metastatic growth in the absence of a primary tumor would provide the opportunity to determine if dendrimer-siRNAs can be delivered directly to secondary metastatic lesions, a property that would benefit patients with advanced, metastatic disease.

Use of siRNA therapeutics targeting nonenzymatic adaptor proteins like IRS2 has the potential to simultaneously modulate signal transduction downstream of several receptors. IRS2 not only serves as an adaptor for IIS receptors, but also for integrin, cytokine, and other growth factor receptors (41,54,55). A single therapeutic that targets IRS2, therefore, is likely to inhibit several parallel processes that contribute to cancer progression, which could provide more clinical benefit than currently available inhibitors that suppress a narrower scope of signaling pathways. The sequence-specific precision of siRNA technology would enable silencing of other adaptors implicated in breast as well as other cancers, such as SHC, GRB2, CRK, and MYD88 (56–59), to inhibit additional downstream signaling pathways as a single or combinatorial interventional approach, especially those for which small molecule inhibitors have not yet been developed.

In summary, our results demonstrate that albumin-binding siRNAs are robustly and efficiently delivered to diverse cell populations within TNBC tumors. In addition, we show that silencing the intracellular adaptor protein IRS2 slows breast tumor growth without causing hyperglycemia, potentially eliminating a significant source of morbidity for cancer patients who would benefit from therapeutics that target the IIS signaling pathway. Increased expression of intracellular adaptors such as IRS2 is associated with poorer survival in patients with other cancers, such as non-small cell lung cancer (NSCLC), cholangiocarcinoma, and pancreatic ductal adenocarcinoma (60–63), and an siRNA-based approach to inhibit this class of proteins could have broader clinical benefit beyond breast cancer. Our work provides evidence that use of albumin-binding siRNAs to inhibit nonenzymatic oncogenes whose protein products cannot be targeted by currently available therapies is an effective strategy to successfully inhibit tumor progression, suggesting that further development of such therapies as antineoplastic agents has the potential to significantly improve clinical outcomes for oncology patients.

## Materials and Methods

### Cell Culture

MDA-MB-231, 4T1, and FL83B cells were obtained from ATCC Cell Biology Collection. PyV-MT mammary tumor cells were isolated from female *FVB MMTV-PyV-MT:Irs1^fl/fl^Irs2^fl/fl^* mice, and PyV-MT*:Irs1^-/-^/Irs2*^-/-^ cells were generated by infection with adenoviral Cre recombinase as previously described (64). MDA-MB-231 and 4T1 cells were grown in RPMI media (Gibco) containing 10% FBS, FL83B cells were grown in F-12K media (Gibco) containing 10% FBS, and PyV-MT cells were grown in DMEM media containing 10%FBS. All cells were grown at 37°C with 5% CO_2_ supply and tested negative for mycoplasma by PCR (Abm).

### Antibodies

Primary antibodies used for immunoblots include rabbit anti-IRS2 (Cell Signaling Technology, Cat# 4502), rabbit anti-IRS1 (Bethyl Laboratories, Cat# A301-158A), mouse anti-actin (Thermo Fisher Scientific, Cat# MA5-11869), mouse anti-tubulin (Cell Signaling Technology Cat#3873), and mouse anti-GAPDH (Santa Cruz Biotechnology, Cat# sc-32233). Anti-vimentin antibodies (Proteintech, clone 6K21, Rosemont, IL, USA) were used for immunohistochemical analysis. Immunofluorescence was performed using a mouse anti-CD31 (Cell Signaling Technology Cat# 77699), rabbit anti-F4/80 (Cell Signaling Technology #70076) and rabbit anti-E-cadherin (Invitrogen PA5-85088). Fluorochrome-labeled antibodies against mouse CD326 (G8.8), CD45 (30-F11), CD206 (C068C2), CD8a (53-6.7), I-A/I-E (M5/114.15.2), CD4 (RM4-5), CD11c (N418), TCRβ (H57-597), and F4/80 (BM8) were purchased from Biolegend and used for flow cytometric analysis.

### Design of human-specific, mouse-specific, and cross-reactive siRNAs targeting IRS2

siRNAs were designed based on algorithm features developed by Shmushkovich *et al*. (33) Briefly, siRNAs were designed based on a 21-nucleotide targeting sequence from human *IRS2* (accession number NM_003749.3) and mouse *Irs2* (accession number NM_001081212.2). Exclusion criteria for sequences included: 1) >56% GC content, 2) single-nucleotide stretches of four or more, or 3) perfect homology to miRNA seeds at position 2-7 of the antisense strand. Sequences were also excluded if position 2-17 of the antisense strand had full complementarity to non-target mRNAs to minimize off-target effects. Cross-species targeting was determined based on perfect homology of positions 2-17 within the target sequence to both species. siRNAs were named based on the position in the target sequence from which they were extracted (e.g. human IRS2 siRNA 5967 was extracted from positions 5967-5988 of NM_003749.3).

### Oligonucleotide synthesis, deprotection, and purification

One set of 48 fully-chemically modified antisense and sense strands was synthesized for *in vitro* screening at a 1μmol scale using standard solid-phase phosphoramidite chemistry on a Dr. Oligo 48 high-throughput oligonucleotide synthesizer (Biolytic, Fremont, CA). Compounds for *in vivo* studies were synthesized on diverse scales using a MerMade 12 synthesizer (Biosearch Technologies, Novato, CA). Standard RNA 2′-O-methyl, 2′-fluoro modifications were applied to improve siRNA stability. Extended nucleic acid (exNA)-modified RNA (exNA) was used at the 3’ terminus of the antisense strands in compounds used for *in vivo* tumor silencing studies (34). Phosphoramidites were purchased from Chemgenes, Wilmington, MA and Hongene Biotech, Union City, CA.

Sense strands of *in vitro* compounds were synthesized on a cholesterol-conjugated solid support (Chemgenes). DCA conjugated sense strands were synthesized on custom-functionalized controlled pore glass (CPG) supports as previously described (26), containing a (dT)_2_ cleavable linker between the oligonucleotide and the conjugate. Synthesis of the amphiphilic dendrimer-conjugated sense strand was carried out on Unylinker CPG (ChemGenes) and commercially available amidites were used to build the dendritic moiety on the 5′ end as previously described (65,66). Cy3-phosphoramidites (GenePharma) were used for fluorescence labeling of the 5’ of sense strands for *in vivo* studies. For Cy3-labeled dendrimer strands, Cy3-DMT-phosphoramidite (Chemgenes) was placed in between the conjugate and the sense strand. Antisense strands were synthesized on Unylinker CPG (Chemgenes), bis-cyanoethyl-N, N-diisopropyl CED phosphoramidite (Chemgenes) was used to introduce a 5’-mono-phosphate for *in vitro* experiments, and 5’-(E)-vinylphosphonate-2’-O-Me-Uridine phosphoramidite (Hongene) was applied for *in vivo* studies.

*In vitro* strands were cleaved and deprotected on synthesis columns (on-column) with ammonia gas (Airgas Specialty Gases) for 90 minutes at 65°C followed by on-column ethanol precipitation and elution with nuclease-free water. *In vivo* sense strands were cleaved and deprotected using 28% aqueous ammonium hydroxide solution for 20 hours at 55°C. *In vivo* antisense strands were cleaved and deprotected using 3% diethylamine in aqueous ammonium hydroxide solution for 20 hours at 35°C, while a 1:1 mixture of ammonium hydroxide and 40% aqueous monomethylamine was used for the sense strands for 120 minutes at 25°C. The controlled pore glass was subsequently filtered and rinsed with 50% ethanol in water and dried overnight under vacuum. Oligonucleotides for *in vivo* experiments were HPLC-purified using an Agilent 1290 Infinity II system (Agilent Technologies, Santa Clara CA) with a PRP-C18 column for Cy3 labeled and lipid-conjugated sense strands and an ion-exchange column for antisense and dendrimer-conjugated sense strands. Purified oligonucleotides were desalted by size-exclusion chromatography and characterized by LC-MS analysis on an Agilent 6530 accurate-mass quadrupole time-of-flight (Q-TOF) LC/MS (Agilent Technologies).

### Identification of IRS2 selective siRNAs

For *in vitro* screening of siRNA candidates, cells were treated with candidate cholesterol-conjugated siRNAs for 72 hours in a 50/50 mixture of complete growth media supplemented with 6% FBS and Opti-MEM (Gibco). Screening concentrations for MDA-MB-231 (0.1875μM), FL83B (1.5μM), and 4T1 (1.5μM) cells served as maximal dose for dose-response assays. Cells were then lysed using lysis mixture (Invitrogen #13228) diluted 1:2 with water containing 0.2mg/mL proteinase K (Invitrogen, #25530–049) and heated at 55 °C for 30 minutes before measuring gene expression.

Human *IRS2* and mouse *Irs2* mRNA levels were determined using the Quantigene 2.0 assay (Affymetrix). Cells were lysed in 250μL homogenizing solution (Invitrogen Cat. QG0517) containing 0.2 mg/mL proteinase K (Invitrogen, #25530–049) and heated at 55°C for 30 minutes. Lysed cells were combined with diluted probe sets (mouse *Irs2;* SB-29780, human *IRS2;* SA-10203; ThermoFisher), added to the capture plate and signal was amplified and detected according to manufacturer’s protocol. Luminescence was detected using a SpectraMax M5 microplate reader and SoftMax Pro 6.3 software (Molecular Devices).

For protein analysis, cells were solubilized at 4°C in lysis buffer (20 mM Tris buffer, pH 7.4, 1% Nonidet P-40, 0.137 M NaCl, 10% glycerol) containing protease inhibitors (Roche) and phosphatase inhibitors (Roche).

### *In vivo* siRNA studies

Mice were housed in a pathogen-free animal facilities at UMass Chan Medical School with 12 hour light/12 hour dark cycle and free access to food and water. Female BALB/c mice (Charles River Laboratories) were used for all experiments and were 7- to 9-weeks old of age at the initiation of each study. For all animal studies, mice were injected with either phosphate buffered saline (PBS control) or fully chemically modified (unconjugated, lipid-conjugated, or dendrimer-conjugated) siRNA that was suspended in 1xPBS for a final siRNA concentration of 40mg/kg (150μL injection volume).

To confirm *in vivo* silencing efficacy of IRS2-targeting siRNAs, naïve mice were subcutaneously injected once with either PBS or 40mg/kg siRNA. Animals were euthanized 2 weeks post-injection, and tissues were collected and either placed in RNAlater and stored at 4°C or flash frozen and stored at -80°C. Two 2 mm punches were collected from tissues stored in RNAlater, placed into QIAGEN Collection Microtubes holding 3 mm tungsten beads and lysed in 300 μL homogenizing solution containing 0.2 mg/mL proteinase K using a QIAGEN TissueLyser II. Samples were heated at 65°C for 30 minutes and then centrifuged at 568x*g* for 5 minutes at room temperature to remove debris. mRNA levels were determined using the Quantigene 2.0 assays (Affymetrix). For protein analysis, frozen tissues were homogenized in RIPA buffer containing protease and phosphatase inhibitors.

To determine the optimal siRNA conjugation strategy for mammary tumor delivery, 4T1 cells (1.6×10^5^) suspended in 30 μL PBS were orthotopically injected into the #4 mammary fat pad of 7- to 9-week-old female BALB/c mice. Animals were palpated daily to detect tumor initiation, and palpable tumors were measured daily with calipers to monitor tumor growth. When tumors reached ∼200 mm^3^ in volume, mice were injected either subcutaneously (150 μL) or intratumorally (50 μL) with either PBS or 40 mg/kg siRNA. Animals were euthanized 48 hours post-injection, and tissues were collected for fluorescence microscopy analysis.

To investigate the role of *Irs2* silencing in mammary tumor progression, 4T1 cells (1×10^5^) were injected into the #2 mammary fat pad and palpable tumors were measured daily with calipers. When tumors reached ∼200 mm^3^ in volume, the mice were randomized into groups and injected subcutaneously with either PBS or 40 mg/kg siRNA, and again 72 hours after the first injection. Mice were euthanized 72 hours after the second injection and tissues were collected to measure tumor and secondary tissue siRNA delivery, gene silencing, and histopathology. Tumor volumes were calculated from caliper measurements using the equation (W^2^*L)/2 to track tumor growth *in vivo*, and terminal tumor volumes were calculated using the equation V=(4/3)π*(L/2)*(W/2)*(D/2)(67,68). Tumor measurements were confirmed by an individual who was blinded to experimental intervention.

### Fluorescence Microscopy

A portion of tumors and secondary tissues were paraffin-embedded and sectioned by the Morphology Research Core Facility at UMass Chan Medical School. Sections were stained with 4′,6-diamidino-2-phenylindole (DAPI) to visualize nuclei and fluorescent images were acquired with a Leica DMi8 inverted microscope (Leica Microsystems) and a Nikon A1 confocal microscope (Nikon). Whole-tissue images were taken using a 10X objective and confocal images were taken using a 60X oil immersion objective. Images were adjusted for equal brightness and contrast to compare the degree of siRNA biodistribution.

### Immunofluorescence Staining

Formalin-fixed, paraffin-embedded tissue sections mounted on glass slides were deparaffinized with xylene and rehydrated using the following series of 5-minute ethanol washes: 100%, 95%, 70%, and 50%, with a final wash with deionized water. An antigen retrieval step was performed using heat and sodium citrate unmasking solution (Cell Signaling Technology #14746S). After antigen retrieval, slides were briefly washed with PBS and placed in blocking solution of PBS containing 0.1%Triton-X 100 (PBST) and 1% bovine serum album (BSA) for 1 hour at room temperature. Slides were then incubated in blocking solution containing an anti-CD31, anti-F4/80, or E-cadherin antibody overnight at 4°C. Sections were washed with PBST, then incubated in blocking solution containing secondary antibodies for 1.5 hours at room temperature. Slides were again washed with PBST, incubated in DAPI diluted in PBS to visualize nuclei, and coverslipped using Prolong Glass mounting medium (ThermoFisher Cat#P36984). Slides were allowed to cure overnight and then imaged with a 20X objective using a Leica DMi8 microscope or a 60X objective using a Nikon A1 confocal microscope. Richardson-Lucy deconvolution was applied to confocal images in Figures 2C and 5F using NIS-Elements Advanced Research software (Nikon). For analysis of tumor vascularization, three representative images were taken per tumor, and blood vessel density and cross-sectional area were quantified and confirmed by three individuals who were blinded to experimental intervention using ImageJ software.

### Flow Cytometry

A portion of collected tumors that were stored in Crytostor CS10 media (Sigma-Aldrich) were dissociated into a single cell suspension using the Miltenyi mouse tumor dissociation kit (Miltenyi Biotec Cat#130-096-730) according to manufacturer’s protocol. Cell suspensions were treated with TruStain FcX PLUS (anti-CD16/32) antibody (Biolegend #156603) for 5min at 4C, then stained with primary antibodies suspended in PBS + 1% FBS supplemented with Brilliant Stain Buffer (BD Biosciences #659611). After washing with PBS, cells were stained with a Fixable Viability Dye (Invitrogen #65-0865-14) and fixed using a formaldehyde-based fixative (Miltenyi Cat# 130-090-477). Samples were run on a Cytek Aurora flow cytometer and analyzed using FlowJo v10 software. Single-stained cell suspensions and UltraComp eBeads Plus (ThermoFisher #01-3333-42) compensation beads were used as compensation controls.

### Stem-loop RT-PCR

Biodistribution levels of siRNA in mammary fat pad, tumor, pancreas, and liver tissue were quantified using Stem-loop RT-PCR (69,70). Tissues were lysed in homogenizing solution (Invitrogen) containing 0.2 mg/mL proteinase K, heated to 65°C for 30 minutes, and spun down to remove debris. Stock tissue supernatants were diluted 1:60 with nuclease-free water for downstream analysis. An 8-point standard curve ranging from 0-100 nM was generated using diluted control tissue supernatants spiked with either unconjugated, lipid-conjugated, or dendrimer-conjugated siRNAs as a template for cDNA synthesis. Standard curve and experimental samples were heated to 95°C for 20 minutes and combined with 1μM RT primers, 10 mM dNTPs (New England Biolabs; N0447L), 10X M-MuLV RT reaction buffer (New England Biolabs; B0253S), and M-MuLV reverse transcriptase (New England Biolabs; M0253L) for cDNA synthesis. Synthesis of cDNA was performed using the following protocol: 16°C for 30 minutes for primer annealing, 42°C for 30 minutes for reverse transcription, and then 70°C for 15 minutes for enzyme inactivation. For the qPCR reaction setup, a master-mix consisting of 5 μL of Power-Up SYBR qPCR Mastermix (Applied Biosystems), 0.05 μL forward primer at 100 μM, 0.05 μL universal reverse primer at 100 μM, and 2.9 μL water was mixed with 2 μL of cDNA template diluted 1:1 with nuclease-free water. The thermal cycling program was set as followed: 2 minutes incubation at 50°C, a 2-minute denaturation at 95°C, then 50 cycles of 10 seconds at 95°C and 20 seconds at 60°C.

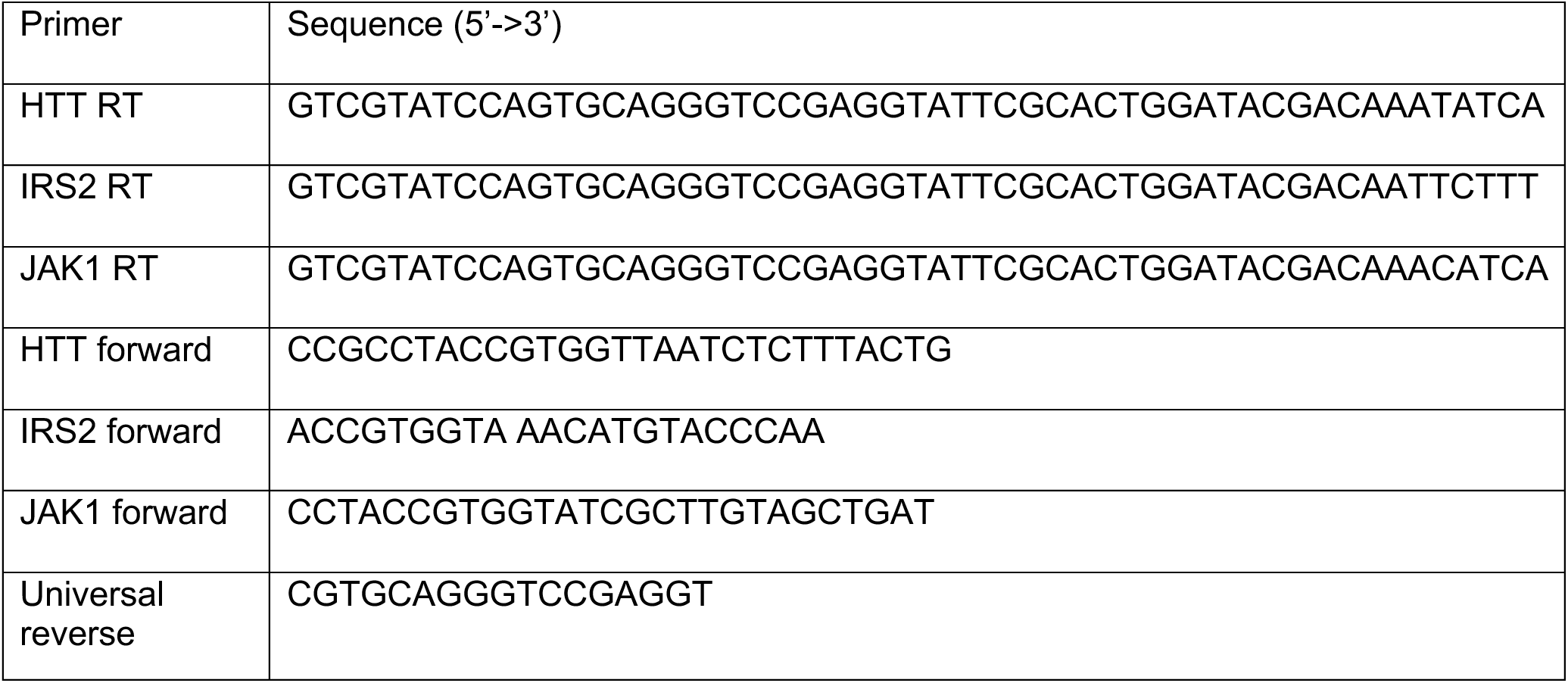

### Serum Glucose Analysis

Whole blood was collected from mice via the facial vein at time of euthanasia and serum was isolated using serum separator tubes (BD #365967). Serum glucose concentrations were then measured using a glucose colorimetric detection kit (ThermoFisher Scientific Cat# EIAGLUC) according to manufacturer’s protocol.

### Immunoblotting

Cell or tissue extracts containing equivalent amounts of protein were resolved by SDS-PAGE and transferred to nitrocellulose membranes. Membranes were blocked for 1 hour at room temperature in 50 mM Tris buffer, pH 7.5 with 0.15 M NaCl and 0.1% Tween-20 (TBST) and 5% (wt/vol) dry milk, incubated overnight at 4°C in TBST containing 3% (wt/vol) bovine serum albumin (BSA) and primary antibodies. Membranes were washed with TBST and then incubated for 1-2 hours at room temperature in 5% blocking buffer with milk containing peroxidase-conjugated secondary antibodies. Bands were detected by chemiluminescence using a ChemiDoc XRS+ system (Biorad) and quantified by densitometry using ImageJ. Signals were normalized to total housekeeping genes to compare relative protein abundance.

### Tumor and Pancreas Histopathology and Immunohistochemistry

Sections of tumors and pancreas tissue were stained with hematoxylin and eosin (H&E) by the Morphology Research Core Facility at UMass Chan Medical School using standard protocols. Stained slides were evaluated for histomorphologic characterization and determination of the percent neoplastic composition of the tissue, percent necrotic composition of the neoplastic population, and the number of mitotic figures per ten contiguous high-powered fields (HPFs; 2.37mm^2^). To determine pancreatic islet cell mass, the total number and cross-sectional area of all islets were compared between experimental groups.

Immunolabeling of FFPE sections of all tumor specimens was performed by using 5 μm sections with a recombinant anti-vimentin or an anti-E-cadherin antibody using standard protocols. Antigen retrieval was performed using sodium citrate solution. Normal murine adipose tissue was used as a positive control while primary antibody was removed in negative controls. Stained specimens were scored as having no (0), weak (1), moderate (2), or strong (3) immunoreactivity, and the percentage of neoplastic cells that stained positive was recorded. All histologic evaluations, including the identification and enumeration of mitotic figures, and immunohistochemical scoring were performed by a board-certified veterinary pathologist who was blinded to experimental intervention.

### Transwell Migration Assays

PyV-MT*: Irs1^-/-^/Irs2*^-/-^ cells stably expressing either a FLAG-tagged IRS2 or an empty vector control were transfected with shRNAs targeting either GFP or vimentin (TRCN0000089828; TRCN0000089830) using Lipofectamine 3000 for 24 hours. Cells were plated into the top chamber of 6.5 mm Transwells (8 μm pore size) (Corning), 3T3-conditioned media was added to the bottom chamber, and cells were allowed to migrate for 5 hours at 37°C. Cells that had migrated to the lower surface of the filters were fixed in methanol for 10 minutes and mounted on glass slides using a Vectashield mounting medium containing DAPI (Vector Laboratories, Burlingame, CA). Migration was quantified by counting the number of stained nuclei in four independent fields in each Transwell.

### Statistical Analysis

All data were analyzed using GraphPad Prism 10 software (GraphPad Software, Inc., San Diego, CA, USA). Student’s t-test was applied and p-value of <0.05 was considered to indicate statistical significance when comparing 2 groups. For comparison of more than 2 groups, one-way ANOVA followed by Tukey’s multiple comparisons was applied.

### Study Approval

All animal experiments were performed in accordance with animal care ethics approval and guidelines of the University of Massachusetts Chan Medical School Institutional Animal Care and Use Committee. All procedures were approved under the Protocol #201900333 (Shaw Laboratory) and #202000010 (Khvorova Laboratory) and in accordance with the National Research Council’s The Guide for the Care and Use of Laboratory Animals. The mice utilized in this study were female because human cancer largely affects female patients.

## Acknowledgments

We thank members of the Shaw, Khvorova, and Mercurio labs for helpful discussion and comments on the manuscript. We thank the University of Massachusetts Chan Morphology Core Facility, particularly Jayme Heywosz and Charissa Le, for assistance in the histological preparations and stains used throughout this project. Illustration figures were created with BioRender.com under the license of UMass Chan Medical School. This work was supported by the National Center for Advancing Translational Sciences grant UL1-TR001453 (to C.E.T. and L.M.S), and National Institutes of Health (NIH) grants R01-CA290778 (to L.M.S.), S10OD20012 and R35-GM131839 (to A.K), and K01-OD034451 (to C.P.). The content is solely the responsibility of the authors and does not necessarily represent the official views of the National Institutes of Health.

## Data and material availability

All data are available in the main text or the supplementary materials. Requests for further information and resources should be directed to and will be fulfilled by LMS. All unique/stable reagents generated in this study are available from LMS with a completed materials transfer agreement.

---

**Supplemental Data Figure 1.**
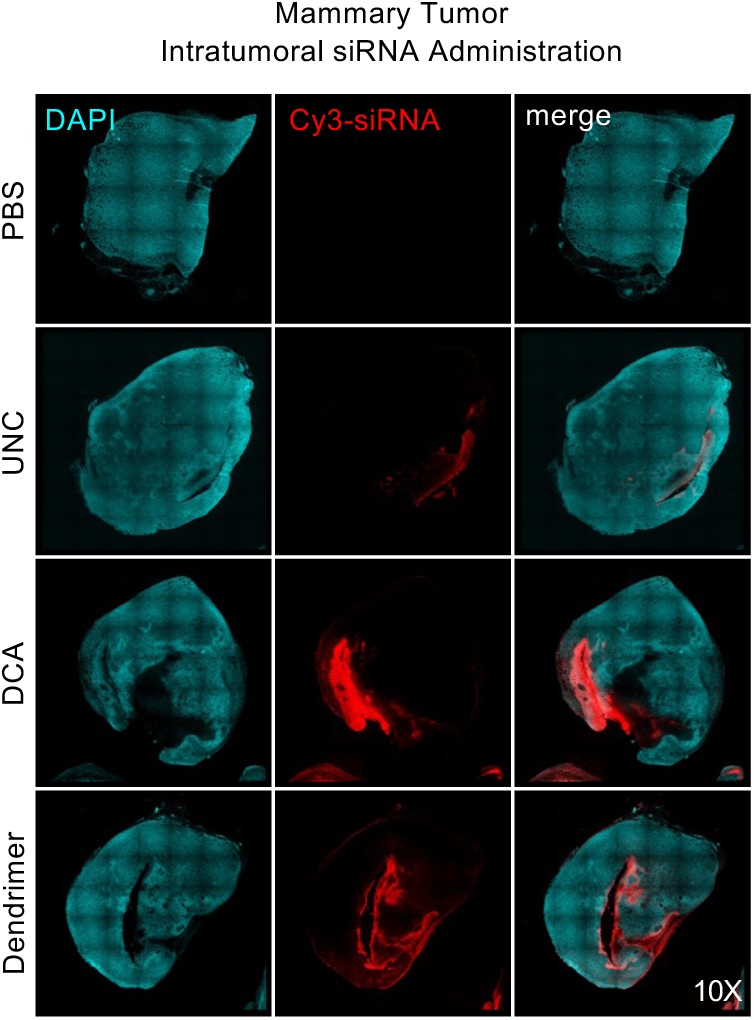
Representative fluorescence microscopy images of mammary tumors 48 hours after intratumoral injection of PBS or 40 mg/kg Cy3-labeled unconjugated (UNC), DCA-conjugated, or dendrimer-conjugated siRNAs. n=5 mice/group.

**Supplemental Data Figure 2.**
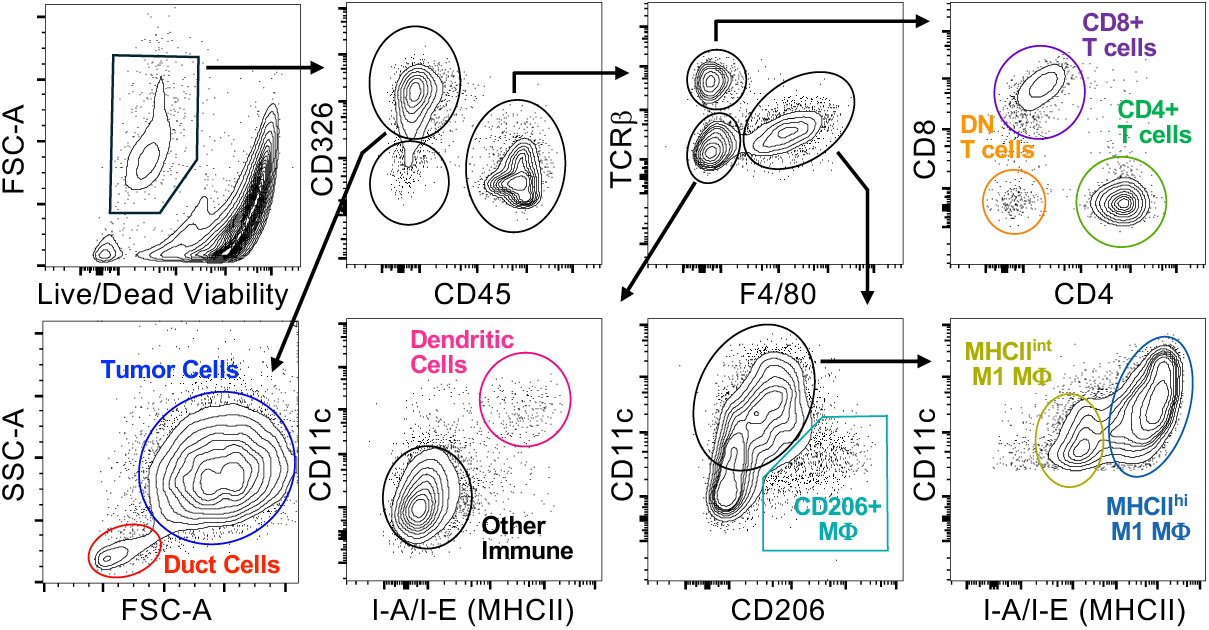
Flow cytometric gating strategy to identify primary tumor cells and tumor stromal and immune cell populations within 4T1 mammary tumors.

**Supplemental Data Figure 3.**
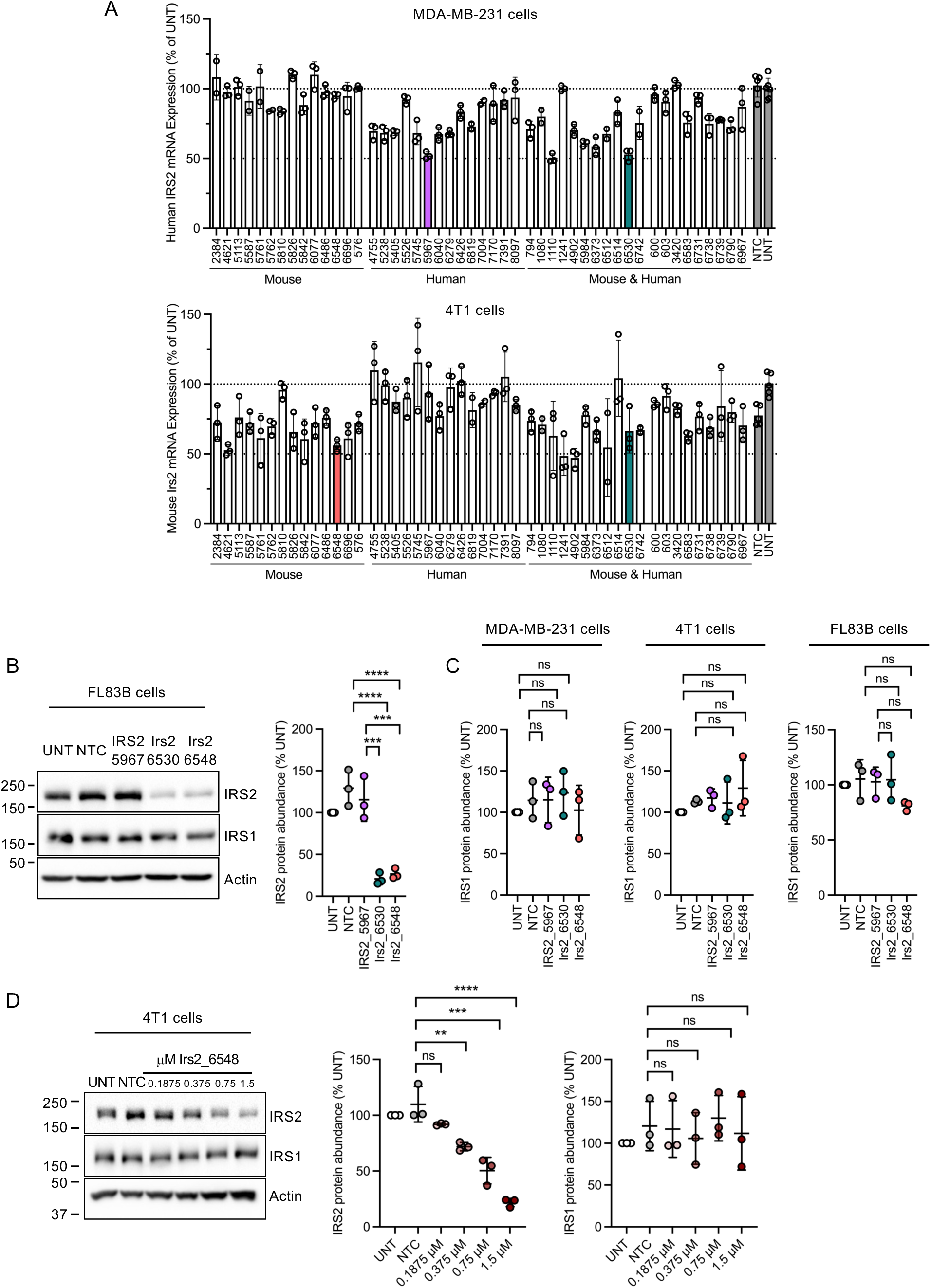
(A) *In vitro* screen of 48 cholesterol-conjugated, fully chemically modified siRNAs designed to silence *IRS2/Irs2* in MDA-MB-231 and 4T1 cells. (B) IRS2 protein abundance after treatment with siRNAs 5967, 6530, and 6548 in the FL83B mouse hepatocyte cell line. (C) Quantification of IRS1 protein abundance in MDA-MB-231, 4T1, and FL83B cells after treatment with siRNAs 5967, 6530, and 6548. (D) IRS2 and IRS1 protein abundance in 4T1 cells after treatment with siRNA 6548 at four doses (0.1875-1.5 μM). NTC: nontargeting control. n=3 independent samples (Supplemental Figure 6A) or 3 independent experiments (Supplemental Figure 6B-D). Data presented as means ±SD. ns nonsignificant **p<0.01 ***p<0.001 ****p<0.0001 by one-way ANOVA.

**Supplemental Data Figure 4.**
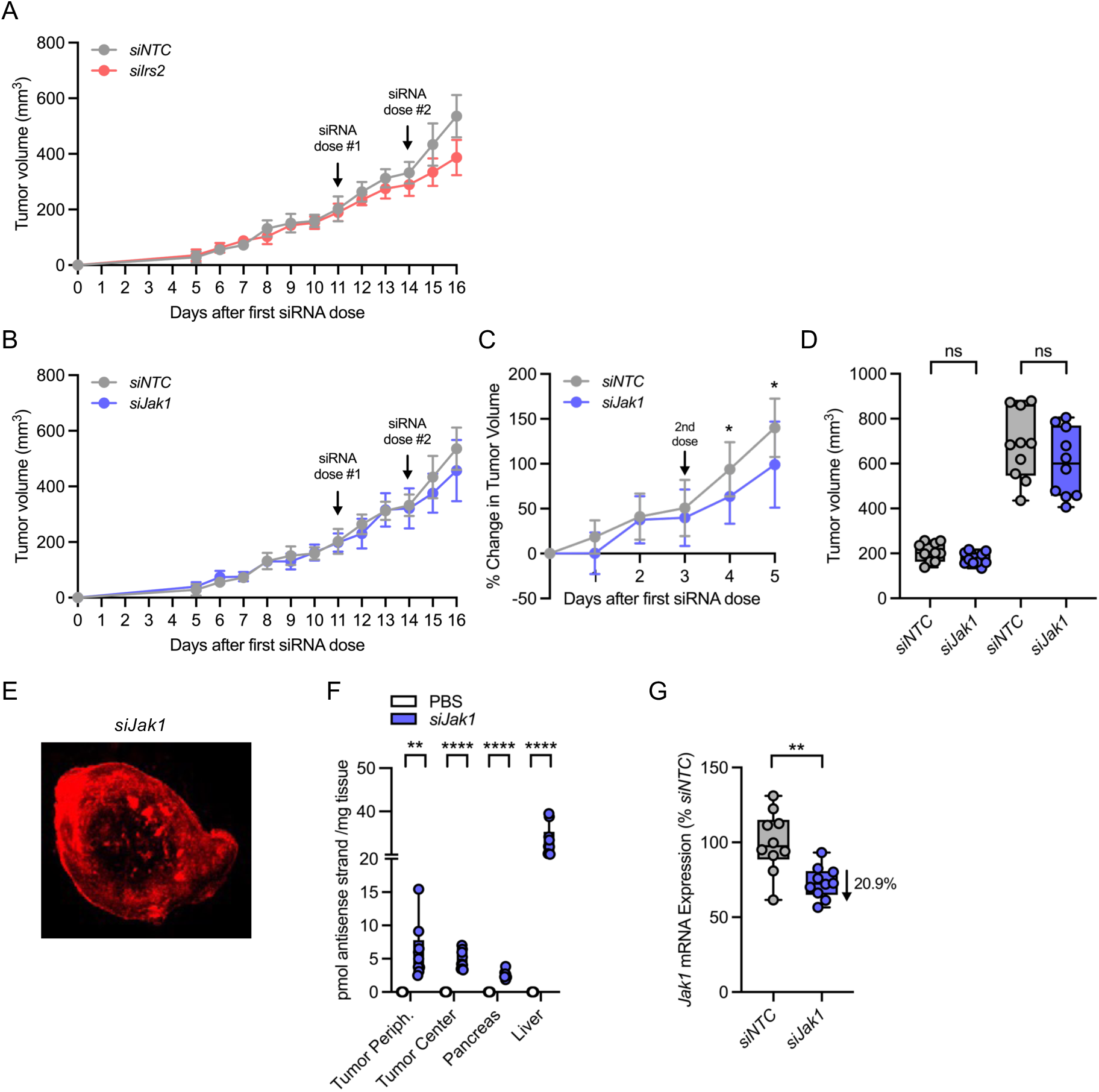
(A-B) 4T1 mammary tumor volume growth curves before and after treatment with *siNTC*, *siIrs2* (A) or *siJak1* (B). (C) 4T1 mammary tumor growth rates after treatment with *siNTC* or *siJak1*. (D) Tumor volumes before *siNTC* or *siJak1* administration and after siRNA treatment. NTC: nontargeting control. Analysis of n=10 tumors. (E) Representative fluorescence microscopy of 4T1 mammary tumor from mice treated with Cy3-labeled *siJak1*. (F) Quantification of Cy3-labeled *siJak1* antisense strand concentrations in 4T1 mammary tumors and secondary tissues. (G) *Jak1* mRNA expression in mammary tumor biopsies from mice treated with *siNTC* or *siJak1*. Line in box-and-whisker plots (Supplemental Figure 4D,F,G) represents median value, while whiskers represent minimum-to-maximum values. Data are presented as means ±SD (Supplemental Figure 4A-C). ns nonsignificant *p<0.05**p<0.01****p<0.0001 by Student’s t-test (Supplemental Figure 4F (by tissue type), 4C (per time point), 4G); ns=nonsignificant by one-way ANOVA (Supplemental Figure 6D).

**Supplemental Data Figure 5.**
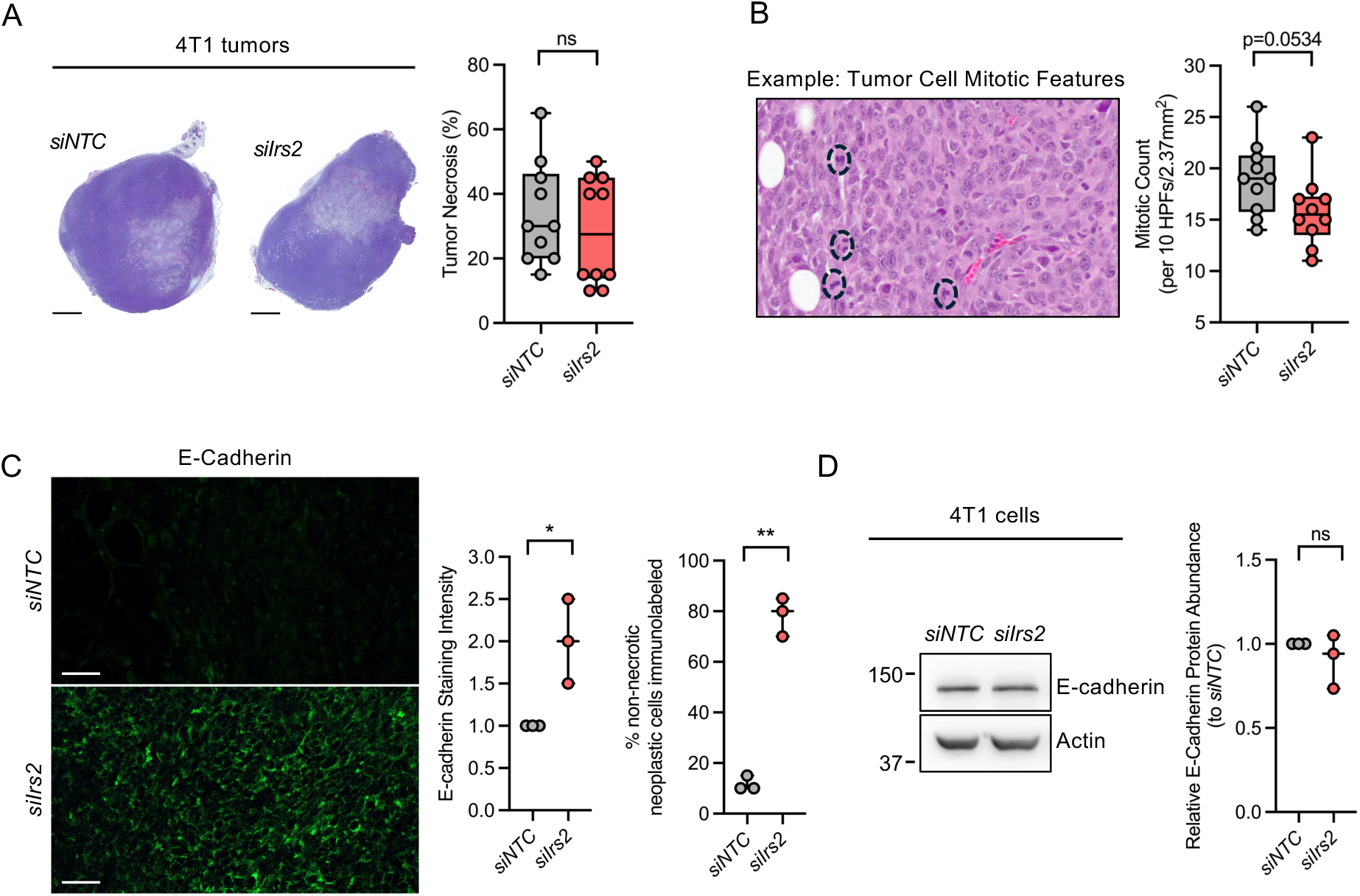
(A) Representative H&E images and quantification of the percent tumor necrosis (right) in 4T1 mammary tumors from mice treated with either *siNTC* or *siIrs2*. (B) Representative H&E image and quantification of mitotic counts in 4T1 mammary tumors from *siNTC* or *siIrs2-*treated mice. HPF: high-powered field. (C) Representative immunofluorescence images and quantification of E-cadherin expression in 4T1 mammary tumors from *siNTC* or *siIrs2*-treated mice. (D) E-cadherin protein abundance in 4T1 cells 72 hours after *in vitro* treatment with cholesterol-conjugated *siNTC* or *siIrs2.* NTC: nontargeting control, Analysis of n=10 tumors (Supplemental Figure 5A-B), 3 tumors (Supplemental Figure 5C) or 3 independent experiments (Supplemental Figure 5D). Scale bar 250 μm (Supplemental Figure 5A) or 50 μm (Supplemental Figure 5B). Line in box-and-whisker plots (Supplemental Figure 5A-B) represents median value, while whiskers represent minimum-to-maximum values. Data are presented as means ± SD (Supplemental Figure 4C-D). ns: nonsignificant *p<0.05**p<0.01 by Student’s t-test.

**Supplemental Data Figure 6.**
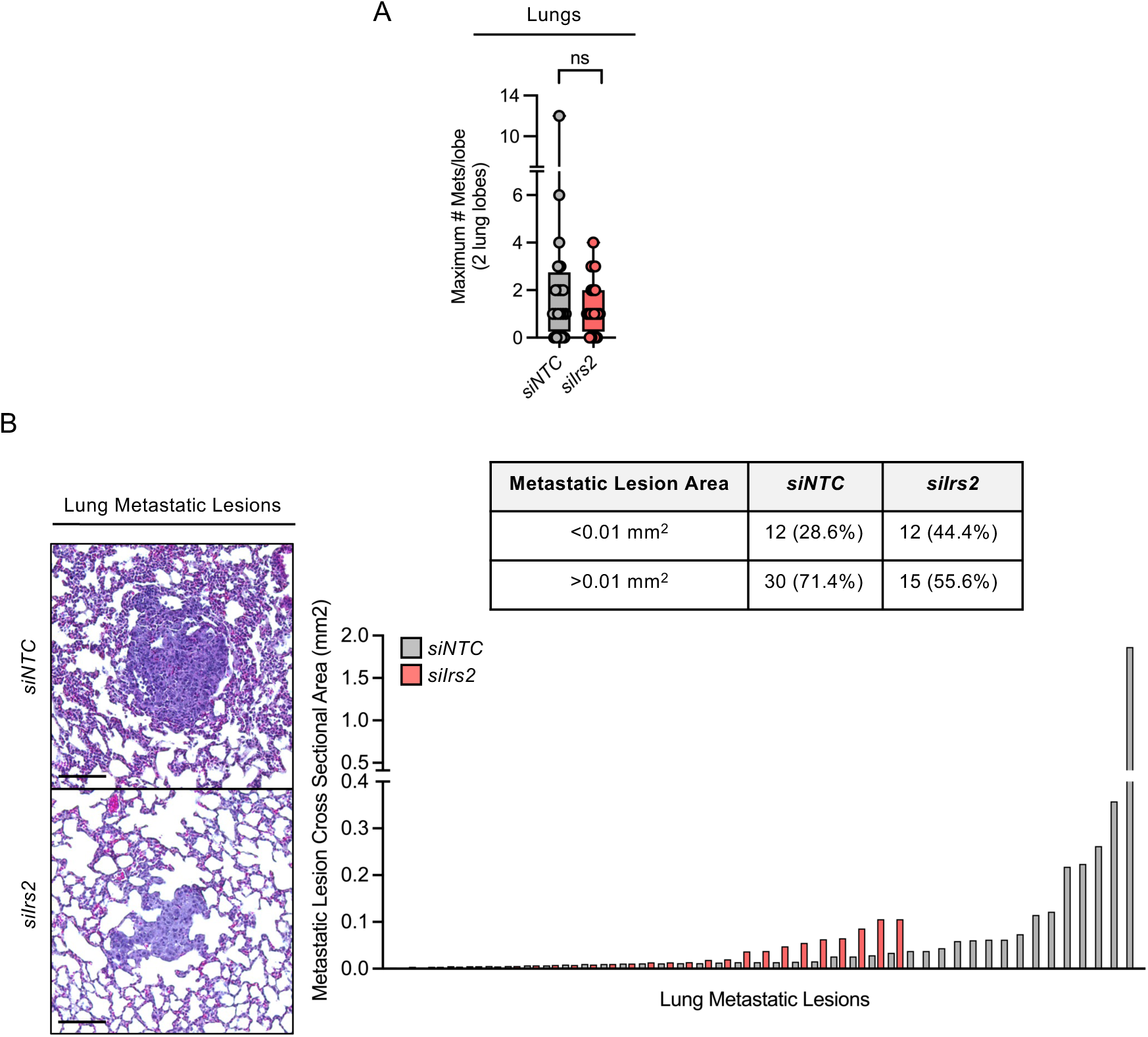
(A) Frequency of metastatic lesions within 2 lung lobes of mice treated with *siNTC* or *siIrs2*. (B) Representative H&E images and quantification of cross-sectional lesion area of metastases within lungs from 4T1 tumor-bearing mice after siRNA treatment. NTC: nontargeting control. Analysis of 2 lung lobes from n=10 mice/group. Scale bar 100 μm. Data are presented as means ±SD. ns nonsignificant by Student’s t-test.

