## Supplemental Table 1 for "Therapeutic delivery of albumin-binding siRNA targeting IRS2 to diverse cell types reduces mammary tumor growth"

**Supplemental Table 1. siRNA sequences and the modification patterns used for *in vitro* screening.**

| | |
|---|---|
| §IRS2_NM_003749_6583_as | ø P(mU)#(fA)#(mA)(fA)(fU)(fA)(mU)(fU)(mG)(fC)(mA)(fU)(mA)(fU)#(mG)#(fC)#(mC)#(mU)#(mU)#(mU)#(fU) |
| IRS2_NM_003749_6731_as | P(mU)#(fA)#(mU)(fA)(fA)(fC)(mU)(fG)(mU)(fU)(mA)(fA)(mA)(fG)#(mU)#(fU)#(mU)#(mA)#(mU)#(mU)#(fU) |
| IRS2_NM_003749_6738_as | P(mU)#(fC)#(mU)(fG)(fU)(fU)(mU)(fA)(mA)(fU)(mA)(fA)(mC)(fU)#(mG)#(fU)#(mU)#(mA)#(mA)#(mU)#(fU) |
| IRS2_NM_003749_6790_as | P(mU)#(fA)#(mU)(fA)(fU)(fA)(mU)(fU)(mU)(fA)(mA)(fC)(mA)(fU)#(mC)#(fU)#(mU)#(mG)#(mA)#(mU)#(fU) |
| IRS2_NM_003749_6967_as | P(mU)#(fA)#(mU)(fU)(fG)(fC)(mA)(fU)(mA)(fU)(mG)(fG)(mC)(fU)#(mA)#(fU)#(mU)#(mA)#(mA)#(mU)#(fU) |
| IRS2_NM_003749_6739_as | P(mU)#(fA)#(mC)(fU)(fG)(fU)(mU)(fU)(mA)(fA)(mU)(fA)(mA)(fC)#(mU)#(fG)#(mU)#(mU)#(mA)#(mU)#(fU) |
| IRS2_NM_003749_600_as | P(mU)#(fU)#(mG)(fU)(fG)(fG)(mU)(fU)(mG)(fU)(mU)(fG)(mU)(fU)#(mG)#(fU)#(mU)#(mG)#(mU)#(mU)#(fU) |
| IRS2_NM_003749_603_as | P(mU)#(fC)#(mG)(fC)(fU)(fG)(mU)(fG)(mG)(fU)(mU)(fG)(mU)(fU)#(mG)#(fU)#(mU)#(mG)#(mU)#(mU)#(fU) |
| IRS2_NM_003749_3420_as | P(mU)#(fG)#(mG)(fC)(fU)(fG)(mU)(fC)(mG)(fC)(mU)(fG)(mC)(fU)#(mG)#(fG)#(mU)#(mG)#(mC)#(mU)#(fU) |
| IRS2_NM_003749_5238_as | P(mU)#(fA)#(mU)(fA)(fA)(fU)(mU)(fA)(mU)(fU)(mA)(fU)(mU)(fU)#(mA)#(fG)#(mU)#(mA)#(mA)#(mU)#(fU) |
| IRS2_NM_003749_6040_as | P(mU)#(fU)#(mG)(fC)(fA)(fU)(mU)(fA)(mU)(fU)(mC)(fU)(mA)(fU)#(mA)#(fG)#(mC)#(mG)#(mA)#(mU)#(fU) |
| IRS2_NM_003749_8097_as | P(mU)#(fC)#(mA)(fC)(fA)(fU)(mU)(fA)(mU)(fU)(mA)(fC)(mA)(fU)#(mG)#(fA)#(mC)#(mU)#(mA)#(mU)#(fU) |
| IRS2_NM_003749_6279_as | P(mU)#(fC)#(mU)(fG)(fA)(fA)(mA)(fU)(mU)(fG)(mA)(fC)(mC)(fU)#(mU)#(fC)#(mC)#(mA)#(mG)#(mU)#(fU) |
| IRS2_NM_003749_7170_as | P(mU)#(fA)#(mA)(fU)(fG)(fU)(mU)(fG)(mU)(fU)(mA)(fU)(mC)(fU)#(mG)#(fG)#(mC)#(mA)#(mG)#(mU)#(fU) |
| IRS2_NM_003749_5405_as | P(mU)#(fA)#(mG)(fG)(fC)(fG)(mU)(fU)(mU)(fG)(mC)(fA)(mA)(fU)#(mG)#(fU)#(mG)#(mA)#(mA)#(mU)#(fU) |
| IRS2_NM_003749_6819_as | P(mU)#(fA)#(mG)(fU)(fC)(fU)(mC)(fU)(mU)(fG)(mU)(fC)(mA)(fU)#(mA)#(fU)#(mC)#(mA)#(mC)#(mU)#(fU) |
| IRS2_NM_003749_5526_as | P(mU)#(fC)#(mA)(fA)(fA)(fU)(mC)(fU)(mG)(fC)(mC)(fA)(mA)(fA)#(mC)#(fC)#(mC)#(mG)#(mG)#(mU)#(fU) |
| IRS2_NM_003749_5745_as | P(mU)#(fC)#(mA)(fA)(fG)(fG)(mG)(fU)(mA)(fG)(mU)(fC)(mU)(fU)#(mG)#(fU)#(mC)#(mC)#(mG)#(mU)#(fU) |
| IRS2_NM_003749_5967_as | P(mU)#(fC)#(mA)(fA)(fC)(fA)(mA)(fC)(mU)(fU)(mA)(fC)(mA)(fU)#(mC)#(fU)#(mC)#(mC)#(mA)#(mU)#(fU) |
| IRS2_NM_003749_7004_as | P(mU)#(fG)#(mG)(fA)(fC)(fU)(mG)(fU)(mA)(fU)(mU)(fC)(mA)(fU)#(mU)#(fU)#(mC)#(mA)#(mU)#(mU)#(fU) |
| IRS2_NM_003749_6426_as | P(mU)#(fU)#(mC)(fA)(fA)(fU)(mA)(fC)(mA)(fC)(mU)(fG)(mC)(fU)#(mU)#(fU)#(mC)#(mA)#(mG)#(mU)#(fU) |
| IRS2_NM_003749_4755_as | P(mU)#(fC)#(mA)(fA)(fA)(fC)(mA)(fG)(mC)(fA)(mC)(fA)(mA)(fU)#(mG)#(fA)#(mU)#(mG)#(mA)#(mU)#(fU) |
| IRS2_NM_003749_7391_as | P(mU)#(fU)#(mG)(fG)(fA)(fC)(mU)(fG)(mU)(fG)(mU)(fG)(mC)(fU)#(mC)#(fU)#(mC)#(mA)#(mC)#(mU)#(fU) |
| Irs2_NM_001081212_6512_as | P(mU)#(fU)#(mU)(fA)(fA)(fA)(mG)(fU)(mU)(fU)(mA)(fU)(mA)(fU)#(mA)#(fG)#(mC)#(mA)#(mA)#(mU)#(fU) |
| Irs2_NM_001081212_6514_as | P(mU)#(fU)#(mG)(fU)(fU)(fA)(mA)(fA)(mG)(fU)(mU)(fU)(mA)(fU)#(mA)#(fU)#(mA)#(mG)#(mC)#(mU)#(fU) |
| Irs2_NM_001081212_6530_as | P(mU)#(fA)#(mA)(fC)(fA)(fC)(mU)(fG)(mU)(fU)(mU)(fA)(mA)(fU)#(mA)#(fA)#(mC)#(mU)#(mG)#(mU)#(fU) |
| Irs2_NM_001081212_6373_as | P(mU)#(fG)#(mU)(fA)(fA)(fA)(mU)(fA)(mU)(fU)(mG)(fC)(mA)(fU)#(mA)#(fU)#(mG)#(mC)#(mC)#(mU)#(fU) |
| Irs2_NM_001081212_6742_as | P(mU)#(fA)#(mU)(fA)(fU)(fG)(mG)(fC)(mU)(fA)(mU)(fU)(mA)(fA)#(mG)#(fG)#(mA)#(mG)#(mG)#(mU)#(fU) |
| Irs2_NM_001081212_5984_as | P(mU)#(fU)#(mU)(fC)(fA)(fA)(mC)(fU)(mU)(fU)(mG)(fU)(mC)(fU)#(mU)#(fG)#(mC)#(mA)#(mA)#(mU)#(fU) |
| Irs2_NM_001081212_794_as | P(mU)#(fC)#(mG)(fC)(fG)(fC)(mU)(fU)(mG)(fU)(mU)(fG)(mA)(fU)#(mG)#(fU)#(mU)#(mC)#(mA)#(mU)#(fU) |
| Irs2_NM_001081212_1110_as | P(mU)#(fA)#(mG)(fG)(fU)(fU)(mC)(fU)(mU)(fG)(mC)(fU)(mC)(fU)#(mG)#(fG)#(mC)#(mC)#(mC)#(mU)#(fU) |
| Irs2_NM_001081212_1080_as | P(mU)#(fU)#(mU)(fC)(fA)(fG)(mG)(fU)(mU)(fC)(mA)(fC)(mC)(fU)#(mG)#(fC)#(mC)#(mA)#(mC)#(mU)#(fU) |
| Irs2_NM_001081212_1241_as | P(mU)#(fU)#(mG)(fA)(fA)(fG)(mA)(fA)(mG)(fA)(mA)(fG)(mC)(fU)#(mG)#(fU)#(mC)#(mC)#(mG)#(mU)#(fU) |
| Irs2_NM_001081212_4902_as | P(mU)#(fG)#(mU)(fA)(fA)(fG)(mC)(fC)(mA)(fG)(mU)(fA)(mA)(fA)#(mU)#(fA)#(mC)#(mC)#(mA)#(mU)#(fU) |
| Irs2_NM_001081212_576_as | P(mU)#(fC)#(mG)(fC)(fA)(fC)(mG)(fC)(mU)(fG)(mU)(fG)(mG)(fU)#(mU)#(fG)#(mU)#(mU)#(mG)#(mU)#(fU) |
| Irs2_NM_001081212_5826_as | P(mU)#(fG)#(mA)(fC)(fA)(fC)(mA)(fU)(mA)(fU)(mA)(fU)(mA)(fU)#(mA)#(fU)#(mA)#(mU)#(mA)#(mU)#(fU) |
| Irs2_NM_001081212_6077_as | P(mU)#(fC)#(mA)(fC)(fA)(fG)(mU)(fG)(mC)(fU)(mA)(fU)(mG)(fU)#(mG)#(fG)#(mC)#(mC)#(mC)#(mU)#(fU) |
| Irs2_NM_001081212_4621_as | P(mU)#(fC)#(mU)(fA)(fA)(fU)(mU)(fC)(mA)(fC)(mA)(fG)(mA)(fA)#(mG)#(fU)#(mC)#(mU)#(mC)#(mU)#(fU) |
| Irs2_NM_001081212_5810_as | P(mU)#(fU)#(mA)(fU)(fA)(fU)(mA)(fU)(mA)(fU)(mA)(fC)(mA)(fU)#(mC)#(fC)#(mC)#(mA)#(mA)#(mU)#(fU) |
| Irs2_NM_001081212_6486_as | P(mU)#(fU)#(mU)(fA)(fA)(fA)(mU)(fU)(mU)(fA)(mA)(fG)(mA)(fU)#(mU)#(fA)#(mA)#(mG)#(mU)#(mU)#(fU) |
| Irs2_NM_001081212_5761_as | P(mU)#(fC)#(mA)(fG)(fC)(fA)(mA)(fC)(mU)(fC)(mA)(fC)(mA)(fU)#(mU)#(fU)#(mC)#(mC)#(mA)#(mU)#(fU) |
| Irs2_NM_001081212_6548_as | P(mU)#(fA)#(mA)(fA)(fC)(fA)(mU)(fG)(mU)(fA)(mC)(fC)(mC)(fA)#(mA)#(fA)#(mG)#(mA)#(mA)#(mU)#(fU) |
| Irs2_NM_001081212_5762_as | P(mU)#(fC)#(mC)(fA)(fG)(fC)(mA)(fA)(mC)(fU)(mC)(fA)(mC)(fA)#(mU)#(fU)#(mU)#(mC)#(mC)#(mU)#(fU) |
| Irs2_NM_001081212_6696_as | P(mU)#(fA)#(mG)(fC)(fA)(fA)(mA)(fC)(mU)(fA)(mA)(fU)(mC)(fU)#(mA)#(fC)#(mU)#(mC)#(mC)#(mU)#(fU) |
| Irs2_NM_001081212_5842_as | P(mU)#(fU)#(mG)(fC)(fA)(fU)(mU)(fA)(mU)(fU)(mC)(fU)(mA)(fU)#(mA)#(fG)#(mU)#(mG)#(mA)#(mU)#(fU) |
| Irs2_NM_001081212_5113_as | P(mU)#(fA)#(mA)(fU)(fC)(fC)(mA)(fU)(mC)(fU)(mU)(fC)(mA)(fA)#(mG)#(fU)#(mC)#(mC)#(mA)#(mU)#(fU) |

| | |
|---|---|
| Irs2_NM_001081212_5587_as | P(mU)#(fA)#(mC)(fA)(fA)(fC)(mA)(fC)(mU)(fG)(mA)(fC)(mA)(fU)#(mU)#(fU)#(mC)#(mC)#(mA)#(mU)#(fU) |
| Irs2_NM_001081212_2384_as | P(mU)#(fA)#(mA)(fC)(fC)(fA)(mA)(fU)(mC)(fU)(mC)(fA)(mA)(fU)#(mG)#(fU)#(mC)#(mU)#(mC)#(mU)#(fU) |
| IRS2_NM_003749_6583_s | ¶(mG)#(mC)#(mA)(fU)(mA)(fU)(mG)(fC)(mA)(fA)(mU)(mA)(mU)(fU)#(mU)#(mA)-TegChol |
| IRS2_NM_003749_6731_s | (mA)#(mA)#(mC)(fU)(mU)(fU)(mA)(fA)(mC)(fA)(mG)(mU)(mU)(fA)#(mU)#(mA)-TegChol |
| IRS2_NM_003749_6738_s | (mA)#(mC)#(mA)(fG)(mU)(fU)(mA)(fU)(mU)(fA)(mA)(mA)(mC)(fA)#(mG)#(mA)-TegChol |
| IRS2_NM_003749_6790_s | (mA)#(mG)#(mA)(fU)(mG)(fU)(mU)(fA)(mA)(fA)(mU)(mA)(mU)(fA)#(mU)#(mA)-TegChol |
| IRS2_NM_003749_6967_s | (mA)#(mU)#(mA)(fG)(mC)(fC)(mA)(fU)(mA)(fU)(mG)(mC)(mA)(fA)#(mU)#(mA)-TegChol |
| IRS2_NM_003749_6739_s | (mC)#(mA)#(mG)(fU)(mU)(fA)(mU)(fU)(mA)(fA)(mA)(mC)(mA)(fG)#(mU)#(mA)-TegChol |
| IRS2_NM_003749_600_s | (mA)#(mC)#(mA)(fA)(mC)(fA)(mA)(fC)(mA)(fA)(mC)(mC)(mA)(fC)#(mA)#(mA)-TegChol |
| IRS2_NM_003749_603_s | (mA)#(mC)#(mA)(fA)(mC)(fA)(mA)(fC)(mC)(fA)(mC)(mA)(mG)(fC)#(mG)#(mA)-TegChol |
| IRS2_NM_003749_3420_s | (mC)#(mC)#(mA)(fG)(mC)(fA)(mG)(fC)(mG)(fA)(mC)(mA)(mG)(fC)#(mC)#(mA)-TegChol |
| IRS2_NM_003749_5238_s | (mC)#(mU)#(mA)(fA)(mA)(fU)(mA)(fA)(mU)(fA)(mA)(mU)(mU)(fA)#(mU)#(mA)-TegChol |
| IRS2_NM_003749_6040_s | (mC)#(mU)#(mA)(fU)(mA)(fG)(mA)(fA)(mU)(fA)(mA)(mU)(mG)(fC)#(mA)#(mA)-TegChol |
| IRS2_NM_003749_8097_s | (mU)#(mC)#(mA)(fU)(mG)(fU)(mA)(fA)(mU)(fA)(mA)(mU)(mG)(fU)#(mG)#(mA)-TegChol |
| IRS2_NM_003749_6279_s | (mG)#(mA)#(mA)(fG)(mG)(fU)(mC)(fA)(mA)(fU)(mU)(mU)(mC)(fA)#(mG)#(mA)-TegChol |
| IRS2_NM_003749_7170_s | (mC)#(mC)#(mA)(fG)(mA)(fU)(mA)(fA)(mC)(fA)(mA)(mC)(mA)(fU)#(mU)#(mA)-TegChol |
| IRS2_NM_003749_5405_s | (mA)#(mC)#(mA)(fU)(mU)(fG)(mC)(fA)(mA)(fA)(mC)(mG)(mC)(fC)#(mU)#(mA)-TegChol |
| IRS2_NM_003749_6819_s | (mA)#(mU)#(mA)(fU)(mG)(fA)(mC)(fA)(mA)(fG)(mA)(mG)(mA)(fC)#(mU)#(mA)-TegChol |
| IRS2_NM_003749_5526_s | (mG)#(mG)#(mU)(fU)(mU)(fG)(mG)(fC)(mA)(fG)(mA)(mU)(mU)(fU)#(mG)#(mA)-TegChol |
| IRS2_NM_003749_5745_s | (mA)#(mC)#(mA)(fA)(mG)(fA)(mC)(fU)(mA)(fC)(mC)(mC)(mU)(fU)#(mG)#(mA)-TegChol |
| IRS2_NM_003749_5967_s | (mA)#(mG)#(mA)(fU)(mG)(fU)(mA)(fA)(mG)(fU)(mU)(mG)(mU)(fU)#(mG)#(mA)-TegChol |
| IRS2_NM_003749_7004_s | (mA)#(mA)#(mA)(fU)(mG)(fA)(mA)(fU)(mA)(fC)(mA)(mG)(mU)(fC)#(mC)#(mA)-TegChol |
| IRS2_NM_003749_6426_s | (mA)#(mA)#(mA)(fG)(mC)(fA)(mG)(fU)(mG)(fU)(mA)(mU)(mU)(fG)#(mA)#(mA)-TegChol |
| IRS2_NM_003749_4755_s | (mU)#(mC)#(mA)(fU)(mU)(fG)(mU)(fG)(mC)(fU)(mG)(mU)(mU)(fU)#(mG)#(mA)-TegChol |
| IRS2_NM_003749_7391_s | (mA)#(mG)#(mA)(fG)(mC)(fA)(mC)(fA)(mC)(fA)(mG)(mU)(mC)(fC)#(mA)#(mA)-TegChol |
| Irs2_NM_001081212_6512_s | (mC)#(mU)#(mA)(fU)(mA)(fU)(mA)(fA)(mA)(fC)(mU)(mU)(mU)(fA)#(mA)#(mA)-TegChol |
| Irs2_NM_001081212_6514_s | (mA)#(mU)#(mA)(fU)(mA)(fA)(mA)(fC)(mU)(fU)(mU)(mA)(mA)(fC)#(mA)#(mA)-TegChol |
| Irs2_NM_001081212_6530_s | (mU)#(mU)#(mA)(fU)(mU)(fA)(mA)(fA)(mC)(fA)(mG)(mU)(mG)(fU)#(mU)#(mA)-TegChol |
| Irs2_NM_001081212_6373_s | (mA)#(mU)#(mA)(fU)(mG)(fC)(mA)(fA)(mU)(fA)(mU)(mU)(mU)(fA)#(mC)#(mA)-TegChol |
| Irs2_NM_001081212_6742_s | (mC)#(mC)#(mU)(fU)(mA)(fA)(mU)(fA)(mG)(fC)(mC)(mA)(mU)(fA)#(mU)#(mA)-TegChol |
| Irs2_NM_001081212_5984_s | (mC)#(mA)#(mA)(fG)(mA)(fC)(mA)(fA)(mA)(fG)(mU)(mU)(mG)(fA)#(mA)#(mA)-TegChol |
| Irs2_NM_001081212_794_s | (mA)#(mC)#(mA)(fU)(mC)(fA)(mA)(fC)(mA)(fA)(mG)(mC)(mG)(fC)#(mG)#(mA)-TegChol |
| Irs2_NM_001081212_1110_s | (mC)#(mC)#(mA)(fG)(mA)(fG)(mC)(fA)(mA)(fG)(mA)(mA)(mC)(fC)#(mU)#(mA)-TegChol |
| Irs2_NM_001081212_1080_s | (mG)#(mC)#(mA)(fG)(mG)(fU)(mG)(fA)(mA)(fC)(mC)(mU)(mG)(fA)#(mA)#(mA)-TegChol |
| Irs2_NM_001081212_1241_s | (mA)#(mC)#(mA)(fG)(mC)(fU)(mU)(fC)(mU)(fU)(mC)(mU)(mU)(fC)#(mA)#(mA)-TegChol |
| Irs2_NM_001081212_4902_s | (mU)#(mA)#(mU)(fU)(mU)(fA)(mC)(fU)(mG)(fG)(mC)(mU)(mU)(fA)#(mC)#(mA)-TegChol |
| Irs2_NM_001081212_576_s | (mC)#(mA)#(mA)(fC)(mC)(fA)(mC)(fA)(mG)(fC)(mG)(mU)(mG)(fC)#(mG)#(mA)-TegChol |
| Irs2_NM_001081212_5826_s | (mA)#(mU)#(mA)(fU)(mA)(fU)(mA)(fU)(mA)(fU)(mG)(mU)(mG)(fU)#(mC)#(mA)-TegChol |
| Irs2_NM_001081212_6077_s | (mC)#(mC)#(mA)(fC)(mA)(fU)(mA)(fG)(mC)(fA)(mC)(mU)(mG)(fU)#(mG)#(mA)-TegChol |
| Irs2_NM_001081212_4621_s | (mA)#(mC)#(mU)(fU)(mC)(fU)(mG)(fU)(mG)(fA)(mA)(mU)(mU)(fA)#(mG)#(mA)-TegChol |
| Irs2_NM_001081212_5810_s | (mG)#(mG)#(mA)(fU)(mG)(fU)(mA)(fU)(mA)(fU)(mA)(mU)(mA)(fU)#(mA)#(mA)-TegChol |
| Irs2_NM_001081212_6486_s | (mU)#(mA)#(mA)(fU)(mC)(fU)(mU)(fA)(mA)(fA)(mU)(mU)(mU)(fA)#(mA)#(mA)-TegChol |
| Irs2_NM_001081212_5761_s | (mA)#(mA)#(mA)(fU)(mG)(fU)(mG)(fA)(mG)(fU)(mU)(mG)(mC)(fU)#(mG)#(mA)-TegChol |
| Irs2_NM_001081212_6548_s | (mU)#(mU)#(mU)(fG)(mG)(fG)(mU)(fA)(mC)(fA)(mU)(mG)(mU)(fU)#(mU)#(mA)-TegChol |
| Irs2_NM_001081212_5762_s | (mA)#(mA)#(mU)(fG)(mU)(fG)(mA)(fG)(mU)(fU)(mG)(mC)(mU)(fG)#(mG)#(mA)-TegChol |
| Irs2_NM_001081212_6696_s | (mG)#(mU)#(mA)(fG)(mA)(fU)(mU)(fA)(mG)(fU)(mU)(mU)(mG)(fC)#(mU)#(mA)-TegChol |
| Irs2_NM_001081212_5842_s | (mC)#(mU)#(mA)(fU)(mA)(fG)(mA)(fA)(mU)(fA)(mA)(mU)(mG)(fC)#(mA)#(mA)-TegChol |
| Irs2_NM_001081212_5113_s | (mA)#(mC)#(mU)(fU)(mG)(fA)(mA)(fG)(mA)(fU)(mG)(mG)(mA)(fU)#(mU)#(mA)-TegChol |

| | |
|---|---|
| Irs2_NM_001081212_5587_s | (mA)#(mA)#(mA)(fU)(mG)(fU)(mC)(fA)(mG)(fU)(mG)(mU)(mU)(fG)#(mU)#(mA)-TegChol |
| Irs2_NM_001081212_2384_s | (mA)#(mC)#(mA)(fU)(mU)(fG)(mA)(fG)(mA)(fU)(mU)(mG)(mG)(fU)#(mU)#(mA)-TegChol |
| NTC_as | P(mU)#(fA)#(mA)(fU)(fC)(fG)(mU)(fA)(mU)(fU)(mU)(fG)(mU)(fC)#(mA)#(fA)#(mU)#(mC)#(mA)#(mU)#(fU) |
| NTC_s | (mU)#(mU)#(mG)(fA)(mC)(fA)(mA)(fA)(mU)(fA)(mC)(mG)(mA)(fU)#(mU)#(mA)-TegChol |

§ Target name_Accession number_mRNA targeting site;
ø siRNA antisense strand sequence;
Modifications: m = 2'-O-methyl; f = 2'-Fluoro; # = Phosphorothioate; P = 5'-Phosphate;
¶ siRNA sense strand sequence and modifications: TegChol = Teg linker + 3'-Cholesterol.
